# Molecular anatomy of eosinophil activation by IL5 and IL33

**DOI:** 10.1101/2022.12.21.521419

**Authors:** Joshua M. Mitchell, Justin W. Mabin, Laura K. Muehlbauer, Douglas S. Annis, Sameer K. Mathur, Mats W. Johansson, Alex S. Hebert, Frances J. Fogerty, Joshua J. Coon, Deane F. Mosher

**Affiliations:** Department of Biomolecular Chemistry, University of Wisconsin-Madison, WI; National Center for Quantitative Biology of Complex Systems, Madison, WI; Department of Chemistry, University of Wisconsin-Madison; Department of Medicine, University of Wisconsin-Madison; Morgridge Institute for Research, Madison, WI

## Abstract

IL5 and IL33 are major activating cytokines that cause circulating eosinophils to polarize, adhere, and release their granule contents. We correlated microscopic features of purified human blood eosinophils stimulated for 10 min with IL5 or IL33 with phosphoproteomic changes determined by multiplexed isobaric labeling. IL5 caused phosphorylation of sites implicated in JAK/STAT signaling and localization of pYSTAT3 to nuclear speckles whereas IL33 caused phosphorylation of sites implicated in NFκB signaling and localization of RELA to nuclear speckles. Phosphosites commonly impacted by IL5 and IL33 were involved in networks associated with cytoskeletal organization and eosinophil adhesion and migration. Many differentially regulated phosphosites were in a diverse set of large proteins—RAB44, a “large RAB” associated with crystalloid granules; NHSL2 and VIM that change localization along with the nucleus during polarization; TNFAIP3 vital for control of NFκB signaling, and SRRM2 and PML that localize, respectively, to nuclear speckles and PML bodies. Gene expression analysis demonstrated differential effects of IL5 and IL33 on IL18, CCL5, CSF1, and TNFSF14. Thus, common effects of IL5 and IL33 on the eosinophil phosphoproteome are important for positioning in tissues, degranulation, and initiation of new protein synthesis whereas specific effects on protein synthesis contribute to phenotypic heterogeneity.

**KEY POINTS:** IL33 and IL5 impact common pathways of eosinophil cytoskeletal reorganization, adhesion and migration.

Each lobe of the human eosinophil nucleus has a specific anatomy poised for new onset of cytokine-specific transcription and splicing.

## INTRODUCTION

Eosinophils are granulocytic white blood cells whose lineage is specified early in hematopoietic development and play major roles in tissue homeostasis and host defense^1,2^. Although a minority of circulating white cells, eosinophils are abundant in bone marrow, gastrointestinal tract, and adipose tissue^3–5^. In addition, mediators (e.g., cytokines) released by other white cells or damaged tissue activate eosinophils to sites of inflammation^6^. Activated eosinophils undergo a morphologic change from an ovoid-to polarized shape, with the eponymous crystalloid granules moving to one end of the cell and the nucleus compacting into a specialized uropod, the nucleopod^6,7^. Polarized eosinophils adhere to and exit blood vessel walls into surrounding tissues^8^. Tissue eosinophils are thought to have homeostatic roles and effector roles that are distinguished by the mix of mediators, oxygen free-radicals, or toxic proteins that are generated^6,9,10^. More studies are needed on how signaling pathways impact eosinophil phenotype and activity, especially in diseases characterized by eosinophil inflammation^11^.

We now compare the molecular and morphologic effects of IL5 and IL33, two cytokines that are involved in allergic diseases such as asthma. IL5 and IL33 play interdependent roles in eosinophil development in bone marrow and maintenance in tissues and in vitro are potent activators of eosinophil shape change, adhesion, and degranulation^1,7,12–15^. IL5 is released by lymphocytes and other white cells and works through JAK/STAT signaling^16,17^. IL33 alerts the immune system to a variety of triggers such as epithelial damage and works through NFκB signaling^18–20^. Humanized monoclonal antibody to IL5 or IL33 each diminish but fail to completely eliminate asthma exacerbations^21–24^. In-depth studies of signaling pathways utilized by IL5 and IL33 may inform the design of better therapeutics that target common features of the signaling pathways^5^. We use a combination of mass spectrometry^25–28^ and confocal microscopy to study eosinophils stimulated by IL33 or IL5. We present deep phosphoproteomic analyses, map signaling networks, and localize a diverse set of large proteins harboring multiple labile phosphorylation sites.

## METHODS

Human blood eosinophils of >98% homogeneity were purified from volunteer subjects with allergic rhinitis or non-severe asthma (**Supplementary Table 1**)^12^. The study was approved by the University of Wisconsin-Madison Health Sciences Institutional Review Board, and informed written consent was obtained. Methods for proteomics and phosphoproteomics with tandem mass tags (TMT)^29^, confocal microscopy^30^, generation of antibodies^30^, and RTqPCR^31^ are described in more detail in the figure legends or supplement.

## RESULTS

### Quantification of eosinophil protein and phosphorylation changes in response to IL5 and IL33

Protein abundance and phosphorylation were compared after 10-min treatments with IL5 or nothing, IL33 or nothing, and IL33 or IL5 using TMT and LC-MS/MS (**Figure 1A**). For each comparison, we split 20 million cells from each of the five donors in half. There were no significant changes in overall protein abundance (**Supplemental Spreadsheet 1**), but significant changes were found in phosphopeptides. Principal component analysis demonstrated clustering of individual donor samples according to treatment(**Figure 1B**). Z-scores of the multiplexed datasets demonstrated that the vast majority of differences among the five donors were within two standard deviations of the mean (**Figure 1C**). These metrics indicate that changes in phosphoproteomic data were driven by cytokine activation rather than donor-to-donor variability.

**Figure 1:**
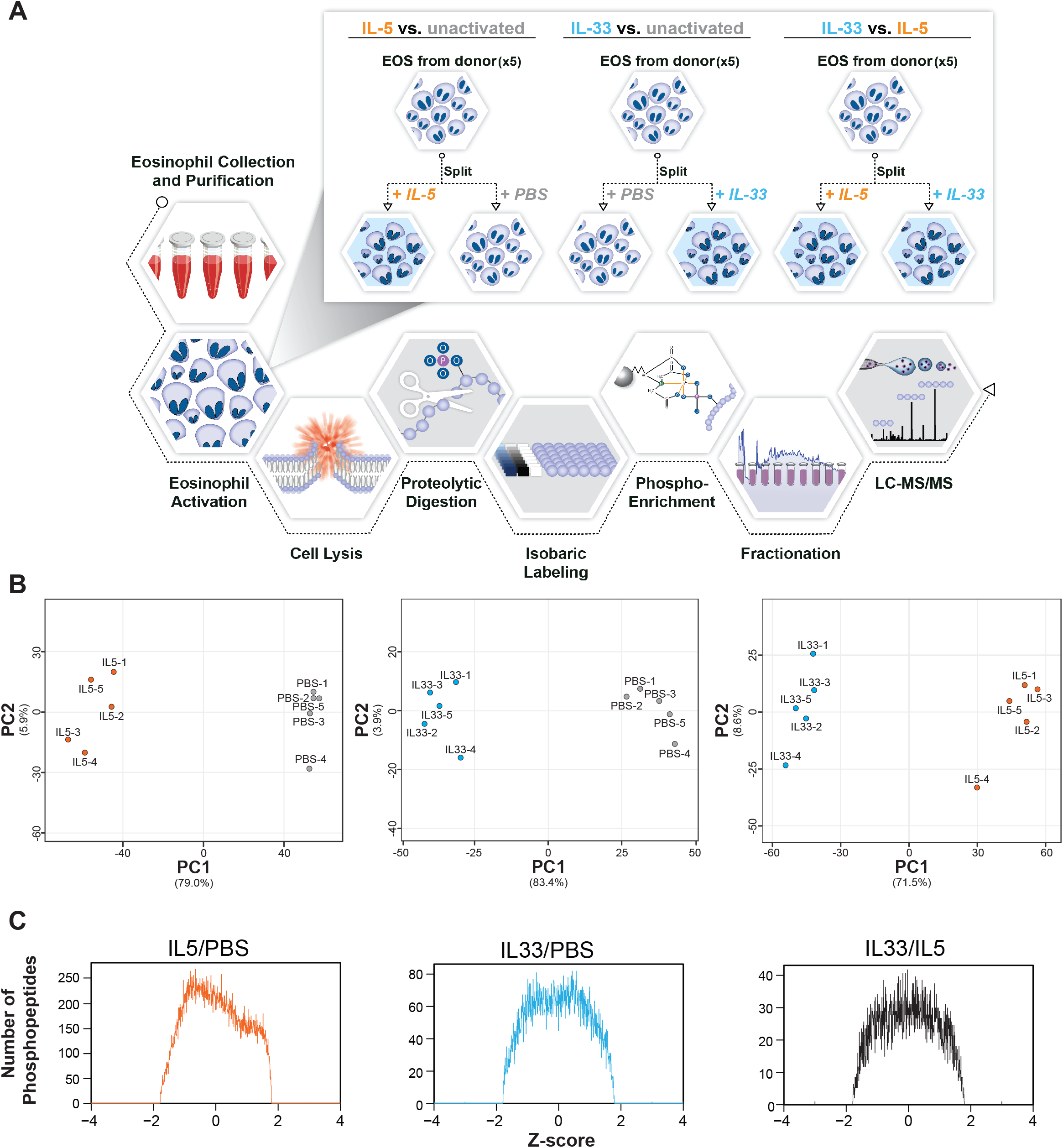
Schematic of the three studies comparing the effects of IL5 and IL33 activation on eosinophil proteomes and phosphoproteomes. **(A)** Schematic of the proteomic and phosphoproteomic methodological approach for the generation of the three compared datasets. **(B)** PCA plots of IL5-versus-non-activated eosinophils, IL33-versus-non-activated eosinophils, and IL33-activated-versus-IL5-activated eosinophils. Unit variance scaling is applied to rows; a singular value decomposition (SVD) with imputation is used to calculate principal components. Axes show principal component 1 and principal component 2 and the percentage of variance explained, respectively. N = 10 data points. **(C)** Frequency histogram of the Z-score differences in phosphopeptides detected in the three phosphoproteomic datasets: IL5-versus-non-activated, IL33-versus-non-activated, and IL33-versus-IL5.

### Phosphoproteomic analysis of IL5 and IL33 activated eosinophils identifies numerous phosphosite changes during cytokine activation

Plots of cytokine-driven differences in individual phosphosites versus relative protein abundance of the proteins harboring the sites (**Figure 2A and 2B**) demonstrated differences in proteins that varied by over orders of magnitude of abundances. Comparing 14,215 phosphosites identified in peptides from IL5-activated versus non-activated eosinophils, 3,940 (28%) in 1,737 proteins were significantly changed (q-value < 0.05, log2 fold change > |0.25|). Of these sites, phosphorylation of 3,063 (78%) increased and of 877 (22%) decreased (**Figure 2A, Supplemental Figure 1A**). Comparing the 4,858 phosphosites identified in IL33-activated versus non-activated eosinophils, 1,695 (35%) in 917 proteins changed significantly, 1,408 (83%) with increased phosphorylation and 287 (17%) with decreased (**Figure 2B, Supplemental Figure 1B**). The plots of foldchange versus abundance revealed many instances of phosphopeptides that share the same horizontal coordinate, indicating that proteins of diverse abundances have multiple significantly changed phosphosites. Indeed, we found that only 34% and 42% of the changed sites in IL5- and IL33-activated eosinophils were in proteins with a single site, whereas others had many, the greatest being NHSL2 with 28 after IL5 treatment (**Figure 2C and 2D**). Remarkably, NHSL2 and RAB44 are highly enriched in proteomes of eosinophils and other granulocytes when compared to other leukocytes^32^.

**Figure 2:**
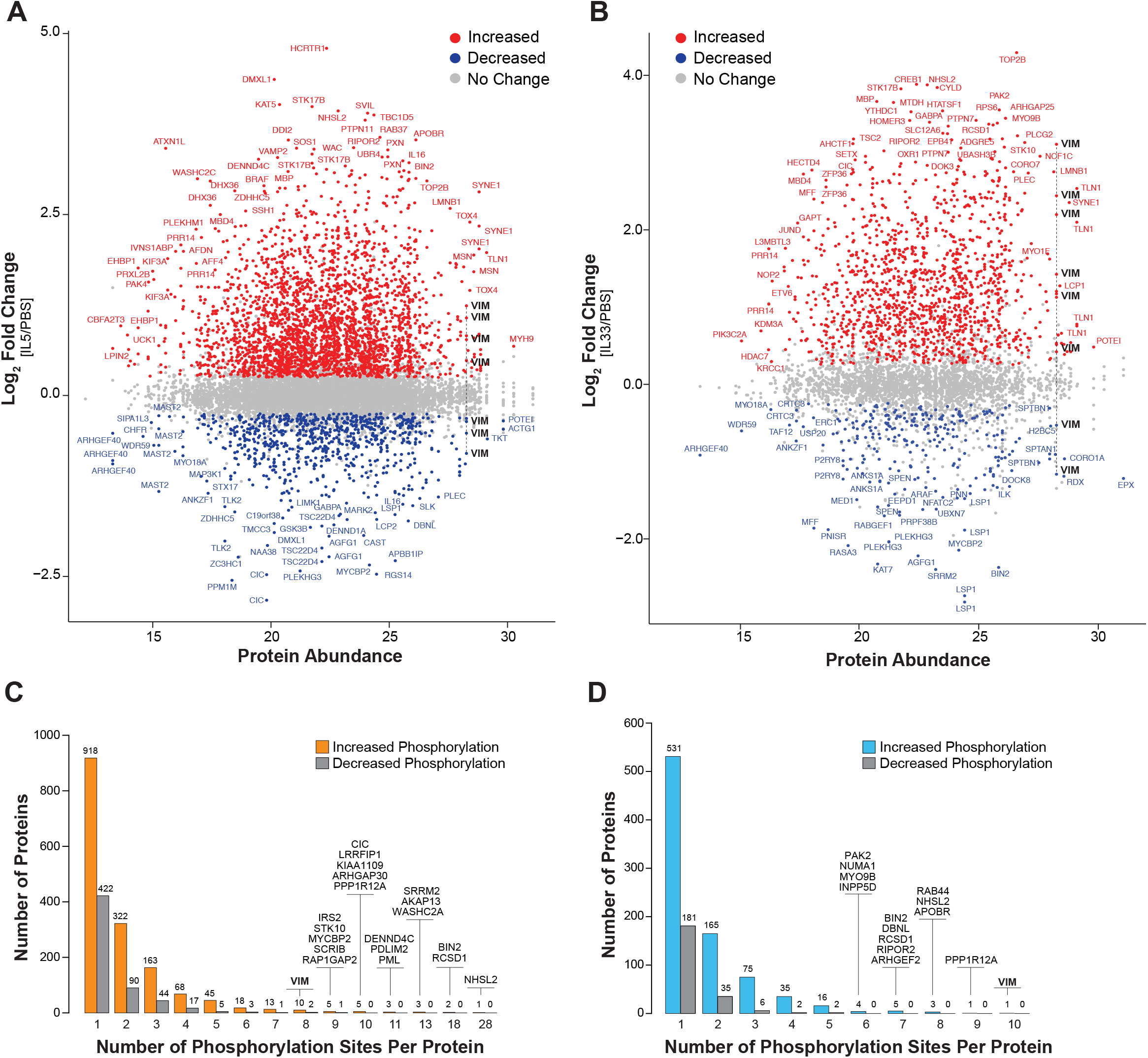
Phosphoproteomic changes of eosinophils treated for 10 min with IL5 or IL33. **(A-B)** MA scatter plots of phosphosites identified in the IL5- **(A)** and IL33- **(B)** versus nonactivated eosinophils. The log2 fold-change of each phosphosite (y-axis) is plotted with respect to its resident protein abundance (x-axis) as previously determined by intensitybased absolute quantification of non-activated eosinophils^64^. Each dot represents one phosphosite. The grey dots indicate no significant change between activated and nonactivated eosinophils, the red represent up-regulated phosphosites and blue dots represent down-regulated phosphosites. VIM is bolded as an example of a protein containing multiple phosphorylation changes. **(C-D)** Bar plots showing the number of proteins (y-axis) with one or several significantly changing phosphosites (x-axis) in IL5- (**C**) or IL33- (**D**) versus non-activated eosinophils. Colored (IL5-orange; IL33-blue) boxes denote increased phosphorylation events, and grey boxes indicate decreasing phosphorylation events. Proteins with the most changing sites are indicated above the corresponding box. VIM is bolded as indicated in A and B.

The IL5-versus-non-activated and IL33-versus-non-activated datasets shared 880 significantly changed phosphosites (**Figure 3A**), which represent 21% (IL5) and 50% (IL33) of total significantly changed phosphosites in the two datasets. Fold-changes of these (**Figure 3B**) correlated with high significance. Most of the phosphosites are found in the lower left and upper right quadrants, indicating changes are in the same direction. Among the sites increased with both cytokines are three in NCF1 for which phosphorylation leads to activation of inducible NADPH oxidase and the oxidative burst (**Figure 3B**)^33^. Multisite phosphorylated proteins shared 144 upregulated and 27 downregulated phosphosites that changed during IL5 and IL33 activation (**Figure 3C**). A heat map of magnitude changes of the 880 phosphopeptides (**Figure 3D**) corroborates the conclusion that the changes are well-coordinated between the activated states, with only 15 of the coincidentally significant sites showing opposite directional change. Proteins with significant phosphosite changes in both IL5- and IL33-activated eosinophils were found in many shared KEGG pathways, including regulation of actin cytoskeleton, focal adhesion and MAPK signaling (**Table 1**). Thus, activation of eosinophils by IL5 and IL33 impact many of the same proteins and pathways.

**Figure 3:**
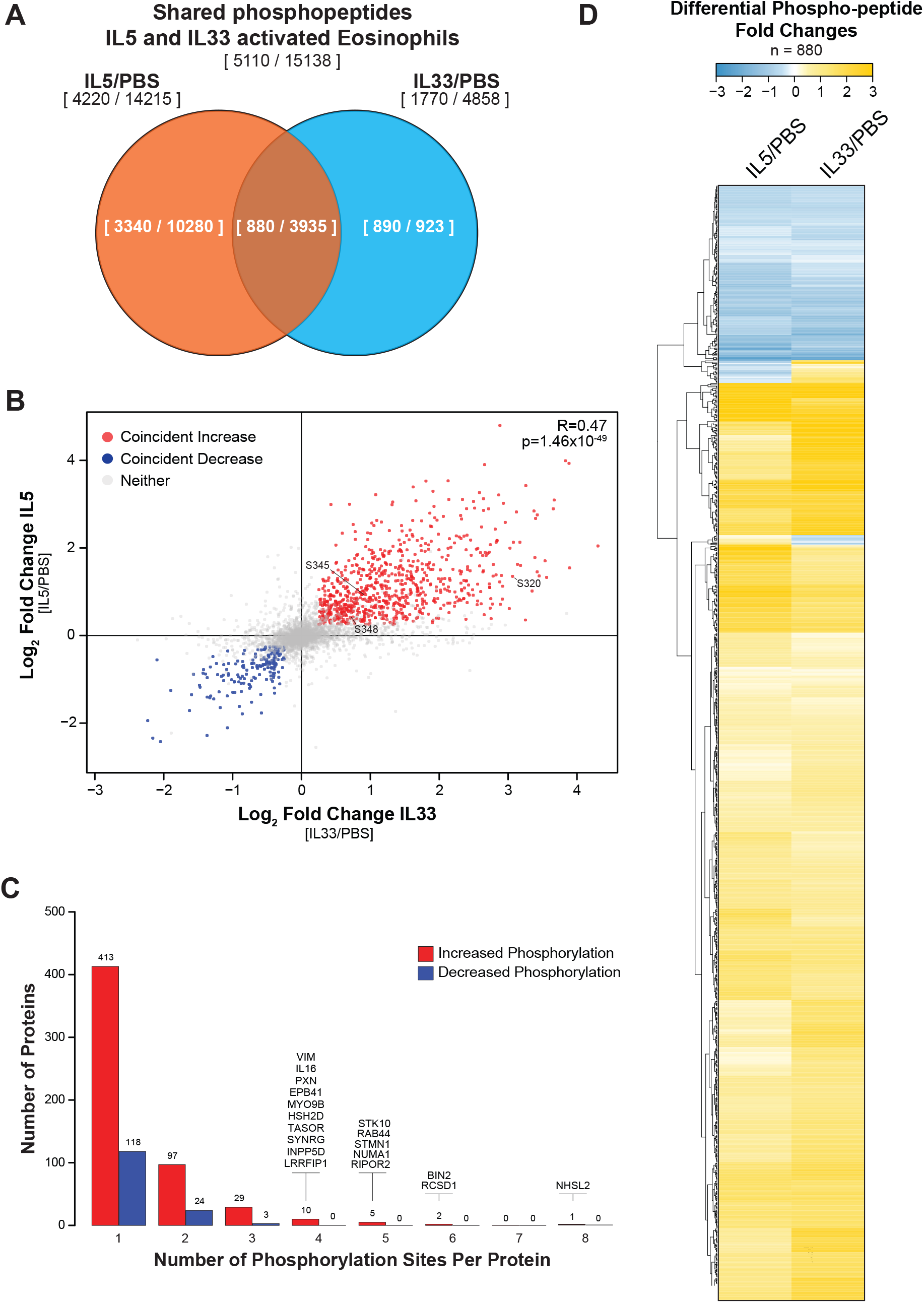
Coincident phosphoproteomic changes of eosinophils treated for 10 min with IL5 or IL33. **(A)** Venn diagram comparing phosphosites found in IL5- and IL33-versus non-activated eosinophils. Ratios show the difference in changed (q < 0.05, log_2_ fold-change > |0.25|) versus total phosphosites identified in the datasets. **(B)** Scatter plot of log_2_ fold changes of phosphosites found in both IL5- (y-axis) and IL33- (x-axis) versus control datasets. The grey dots indicate no significant change between activated states, the red represent coincident up-regulated (right) phosphosites and blue dots represent coincident down-regulated (left) phosphosites. The correlation coefficient and p-value of the comparison of the two treatments is indicated in the top right. Phosphorylations in NCF1 that lead to activation of inducible NADPH oxidase are numbered. **(C)** Bar plot showing the number of proteins (y-axis) with one or several coincidently changing phosphosites (x-axis) in IL5- and IL33-versus non-activated eosinophils. Box color corresponds with the coincident changes in phosphosites described in B. Proteins with the most changing sites are indicated above the corresponding box. **(D)** Heatmap of log2 fold changes of shared phosphosites. Fold change differences in up- and down-regulated phosphosites in yellow and blue, respectively. The hierarchical clustering dendrogram on the left shows the clusters of similarly changing phosphosites between IL5 and IL33 datasets (top).

**Table 1:**
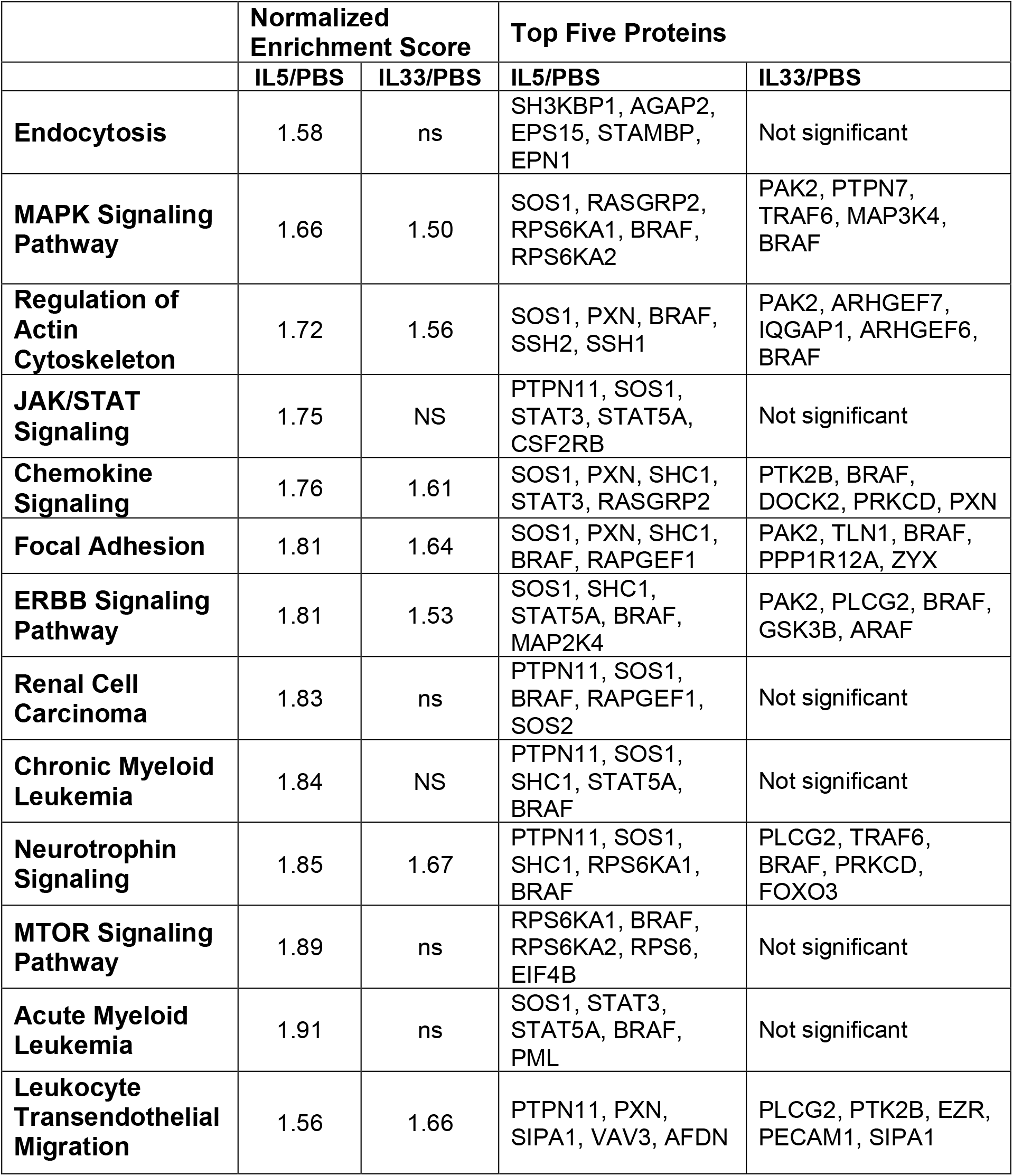
Enrichment of KEGG pathways by proteins harboring sites of increased phosphorylation in the IL5-versus-no treatment (NT) and IL33-versus-NT datasets. NES, normalized enrichment score. Only NES scores that were statistically significant were reported (ns p > 0.05).

### Direct comparison of IL5-versus IL33-activated eosinophils distinguishes the differences in signaling pathway specificity

Comparing fold-changes in cytokine to no-cytokine control datasets is problematic because of the likelihood of variable blunting of differences in intensities among reader ions between the two analyses, known as ratio compression^34^, which underestimates fold-differences. We therefore directly compared eosinophils activated with IL5 or IL33. Of 16,691 identified phosphosites, 4,362 (26%) were significantly different (**Figure 4A** and **Supplemental Figure 2A**). Of these, 2,734 (63%) were more phosphorylated in eosinophils activated by IL33 and 1,628 (37%) in eosinophils activated by IL5 (**Supplemental Figure 2B**). As with the other two datasets, we found proteins that underwent differential multisite phosphorylation events (**Figure 4A and 4B**). Multiple sites were more phosphorylated in VIM, APOBR, CYLD, MYO9B, and TNFAIP3 of cells treated with IL33 whereas the opposite was true for PTPN11, SHC1, SOS1, BIN2, and NHSL2 (**Figures 4A and 4B**).

**Figure 4:**
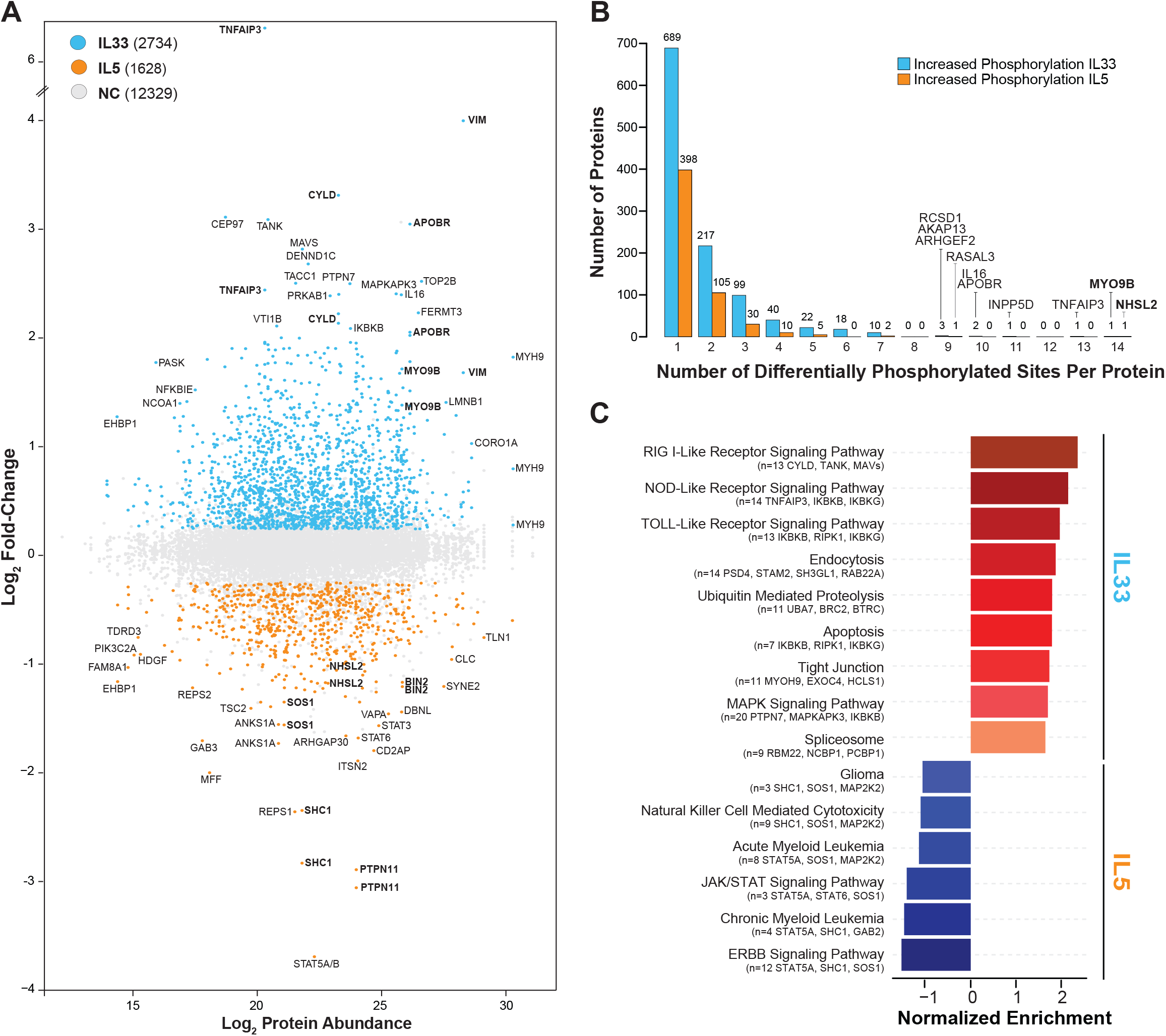
Phosphoproteomic differences of eosinophils treated for 10 min with IL5 or IL33. (A) MA plot of phosphosites identified in the IL33-versus-IL5 dataset. The log2 foldchange of each phosphosite (y-axis) is plotted with respect to its resident protein abundance (x-axis) as described in Figure 2A-B. Each dot represents one phosphosite. The grey dots indicate no significant change between both activated states, the blue dots represent IL33 up-regulated phosphosites and orange dots represent IL5 up-regulated phosphosites. Bolded proteins are discussed in the text. (B) Bar plot showing the number of proteins (y-axis) with one or several significantly changing phosphosites (x-axis) in IL5-versus-IL33 treatment. Colored (IL5-orange; IL33-blue) boxes denote increased phosphorylation events enriched in either activation. Proteins with the most changing sites are indicated above the corresponding box. Bolded MYO9B and NHSL2 are referenced in the text. (C) Gene set enrichment analysis showing the normalized enrichment scores (NES) for the KEGG pathways (left) enriched in IL5 (blue) versus IL33 (red) phosphosites. The total number and list of top proteins are denoted for each KEGG pathway.

Gene set enrichment analysis of KEGG pathways involving the comparative dataset yielded a more focused readout than in the cytokine-versus-nothing datasets (**Figure 4C**). Proteins with more phosphorylation in IL33-treated cells were enriched in RIG I-like, NOD-like, and Toll-like receptor signaling pathways that use the NFκB pathway, whereas those with more phosphorylation in IL5-treated cells were enriched in ERBB and JAK/STAT signaling pathways.

Proteomic and phosphoproteomic datasets were mapped onto templates for IL5 and IL33 signaling pathway (**Figure 5)** using the KEGG Database, WikiPathways, and published networks^35–39^. We first identified eosinophil-expressed proteins identified in at least one of the three datasets (**Supplemental Spreadsheet 1**) and then mapped phosphorylated proteins and significantly changing sites in the IL33-versus-IL5 phosphoproteomic dataset to pathways of interest (**Figure 5**). Finally, we reviewed the map for gaps that could be filled in from the datasets for the individual cytokines versus no treatment.

**Fig. 5:**
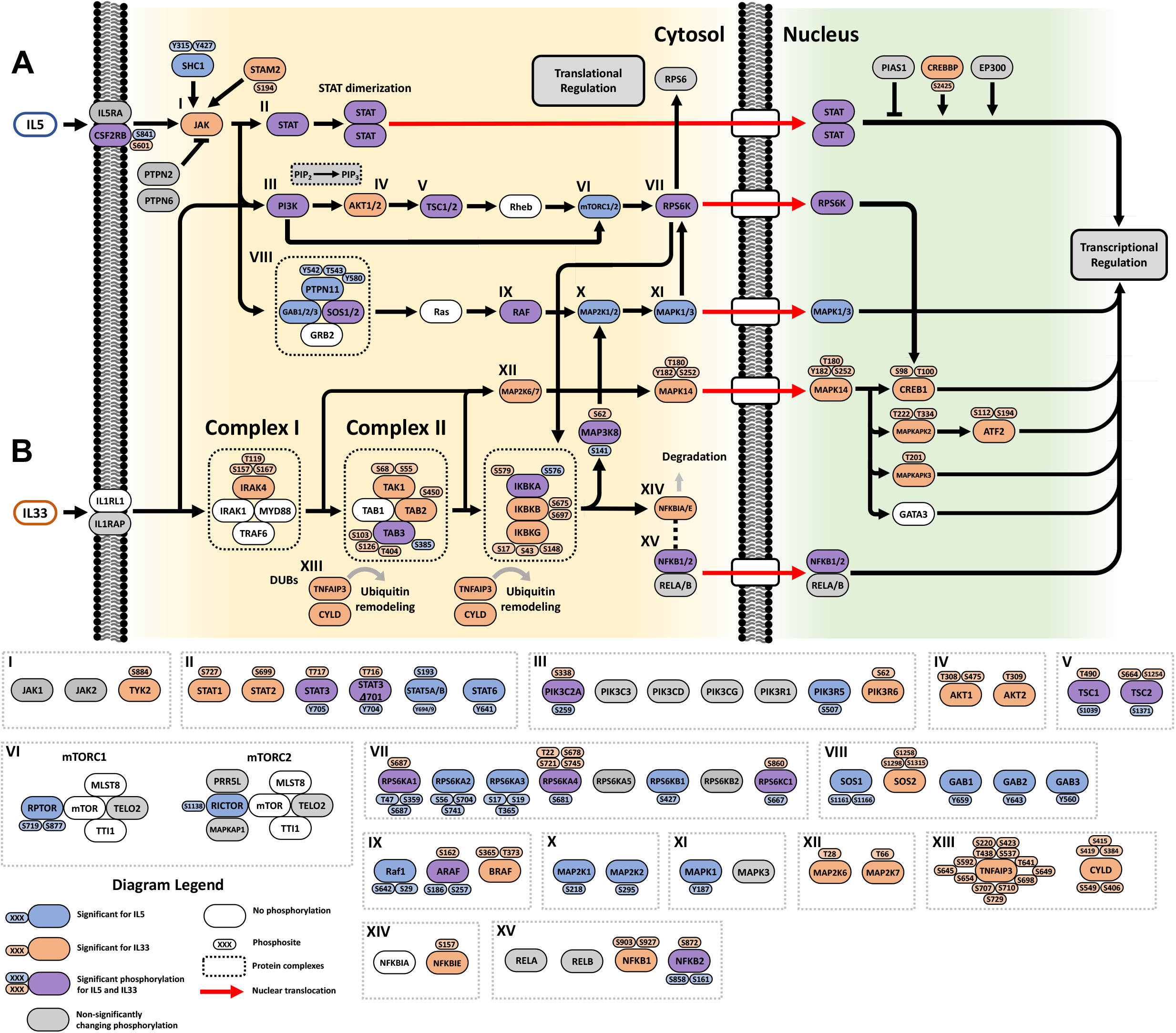
Phosphorylation of signaling pathway proteins upon cytokine exposure in eosinophils. (A) Representative proteins in the IL5—JAK/STAT signaling pathway. (B) Representative proteins in the IL33—NFκB signaling pathway. Nodes (gene names) are colored based upon phosphorylation significance and specificity: white, found proteins for which no phosphosites were found; grey; proteins that are phosphorylated but with no significant differences in sites between IL5- and IL33-activated eosinophils; orange, significantly increased phosphorylation after IL33-activation; blue, significantly increased phosphorylation after IL5-activation; purple, significant increases in phosphorylation of different sites after IL5- and IL33-activation. Phosphosites (residue and position number) are labeled with the indicated color schemes. For protein families that have multiple members and large complexes, the individual members are labeled in the dashed boxes with Roman numerals and are described further in the text. The labeling scheme is summarized at the bottom left.

IL5-induced changes map onto the JAK/STAT pathway (**Figure 5A**). IL5-stimulation causes activating phosphorylation of full-length STAT3 (Y705) and the alternatively spliced ΔS701 proteoform (pY704), as well as of STAT5A/B (Y694/Y699) and STAT6 (Y641) (**Figure 5-II**). These tyrosine phosphorylations allow for homo- and hetero-dimerization and STAT complex translocation into the nucleus^40^. In contrast, IL33 favors phosphorylation of a homologous serine in the more C-terminal transactivation domain (S727 in STAT1) that stabilizes STATs in the nucleus^41,42^.

IL33 signals through the IL1-family and NFκB signaling pathways (**Figure 5B**). The IRAK receptor complex (*Complex I*), Tak1-TAB complex (*Complex II*), and IKK complex harbor numerous labile sites. IKBKG (S17) phosphorylation is required for full transcriptional activation of NFκB^43^. We find evidence for NFκB signaling (**Figure 5-XIV/XV**) with phosphorylation of NFKB1/2. NFKB1 (S903) phosphorylation is permissive for degradation, which can reduce NFKB1 homodimers and promote NFKB response^44,45^. NFKB2 (S872) phosphorylation is important for processing to its active form for transcriptional regulation^46,47^.

PI3K-AKT signaling shows protein phosphosite specificity for each cytokine. PIK3C2A is phosphorylated in both activated states, PIK3R5 is preferential for IL5, and PIK3R6 is preferential for IL33 (**Figure 5-III**). Following PI3K activation, AKT1/2 (T309, T308) phosphorylation activates these kinases (**Figure 5-IV**)^48,49^, leading to TSC2 (S1254) phosphorylation that increases mTORC1 signaling (**Figure 5-V/VI**) ^50^. mTORC1/2 phosphorylation was only apparent in IL5-activated eosinophils. RPTOR phosphorylation appears paradoxical, with S719 upregulating mTORC1 activity and S877 decreasing mTORC1 activity, suggesting a dynamic regulation^51–53^. The RPS6K family of kinases also shows distinct phosphosite specificities (**Figure 5-VII**). IL5 activation upregulates several sites in RPS6KA1/2/3 and B1, while IL33 activation upregulates phosphosites in RPS6KA4.

The SOS-GRB-GAB ternary signaling complex is highly phosphorylated during IL5 activation, which leads to MAPK1 activation (**Figure 5-VIII**). PTPN11 (Y542, Y580) and GAB1/2/3 (Y659/Y643/Y560) tyrosine phosphorylation initiates sequential RAS activation, RAF (S257) activation^54^, MAP2K1/2 (S218) activation^55^, and MAPK1 activation (**Figure 5-IX/X/XI**)^56,57^. SOS1 (S1161) phosphorylation is known to disrupt 14-3-3 protein binding and decreases MAPK activation^58^. In IL33 activation, we see inhibition of this portion of the MAPK pathway with BRAF (S365) phosphorylation maintaining autoinhibition (**Figure 5-IX**)^59^. However, we see alternate IL33 MAPK signaling associated with *Complex I* and *Complex II* with upregulation of MAP2K6 (T28) and MAP2K7 (T66) phosphorylation (**Figure 5-XII**). MAPK14 (T180, Y182) phosphorylation directs its nuclear translocation and subsequent phosphorylation of ATF2, activating target gene transcription^60^. We note significant phosphorylation of nuclear-resident transcription factors CREB1 (S98, T100) and transcription factor kinases MAPKAPK2 (T222, T334), and MAPKAPK3 (T201), which are required for MAPKAPK2/3 activation and nuclear export^61–63^.

Many of the labile phosphosites currently are not known to play a role in activation or inhibition. SOS1/2, PI3K, and RPS6K phosphorylations serve as prime examples of differential regulation of phosphorylation for which we do not know the significance of the changes.

### Focused analysis of proteins with multiple labile phosphorylation sites

Protein phosphorylation state of labile sites is a play-off between kinase and phosphatase activities and location of substrate vis-à-vis location of enzymes. The published eosinophil global proteome reports abundances of 201 of 518 known protein kinases (**Supplemental Figure 3**)^64^. NetPhorest, a web-server algorithm, predicts kinase-substrate relationships based on peptide sequence motifs and known STRING interactions^65^. We used NetPhorest to identify kinases likely to act on the multiple labile sites of, TNFAIP3, PML, RAB44, VIM, NHSL2, and SRRM2 (**Figure 6**) and in parallel studies localized the six proteins in non-activated and activate eosinophils (**Figure 7**) with the goal of also identifying localities of kinase action.

**Figure 6:**
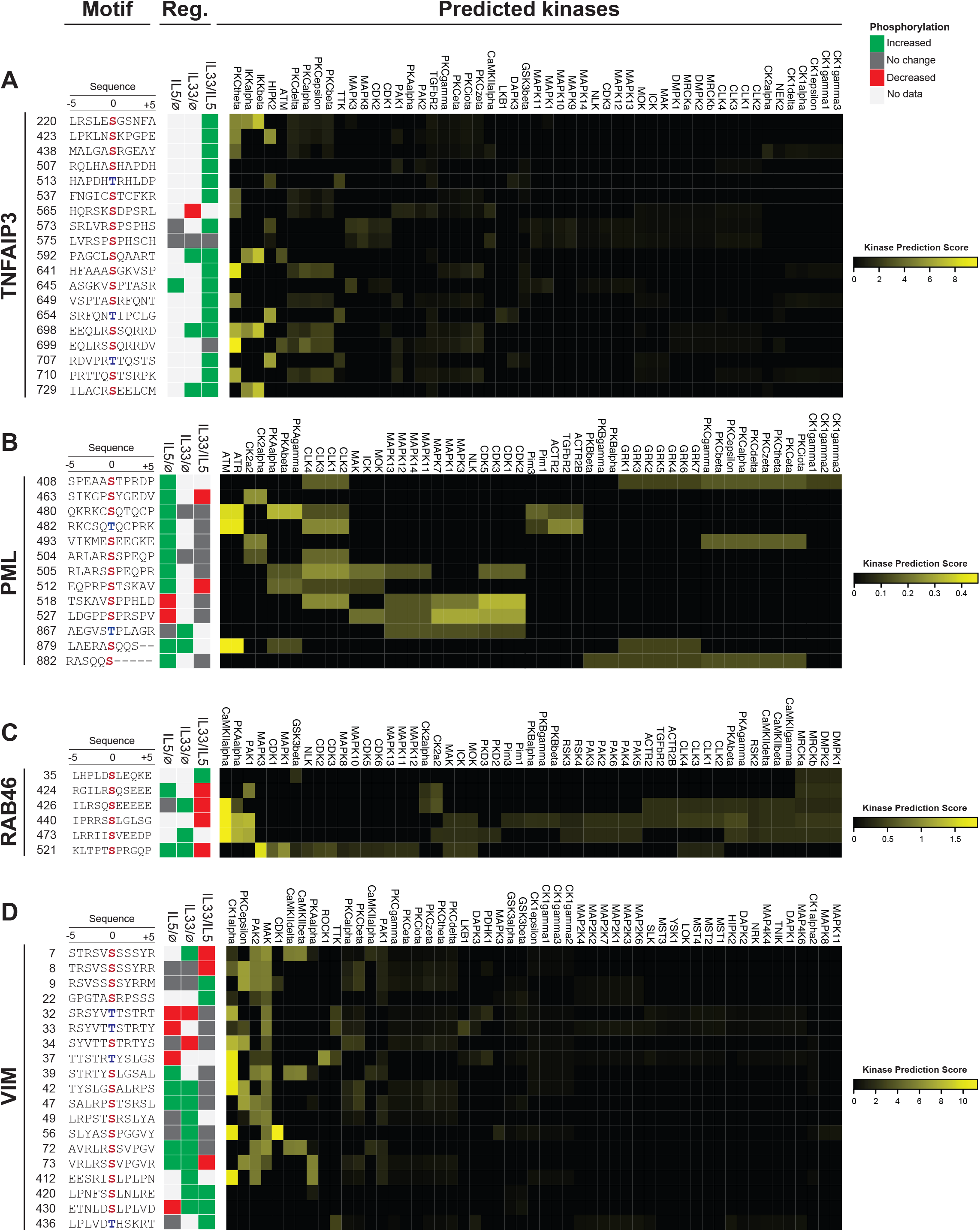
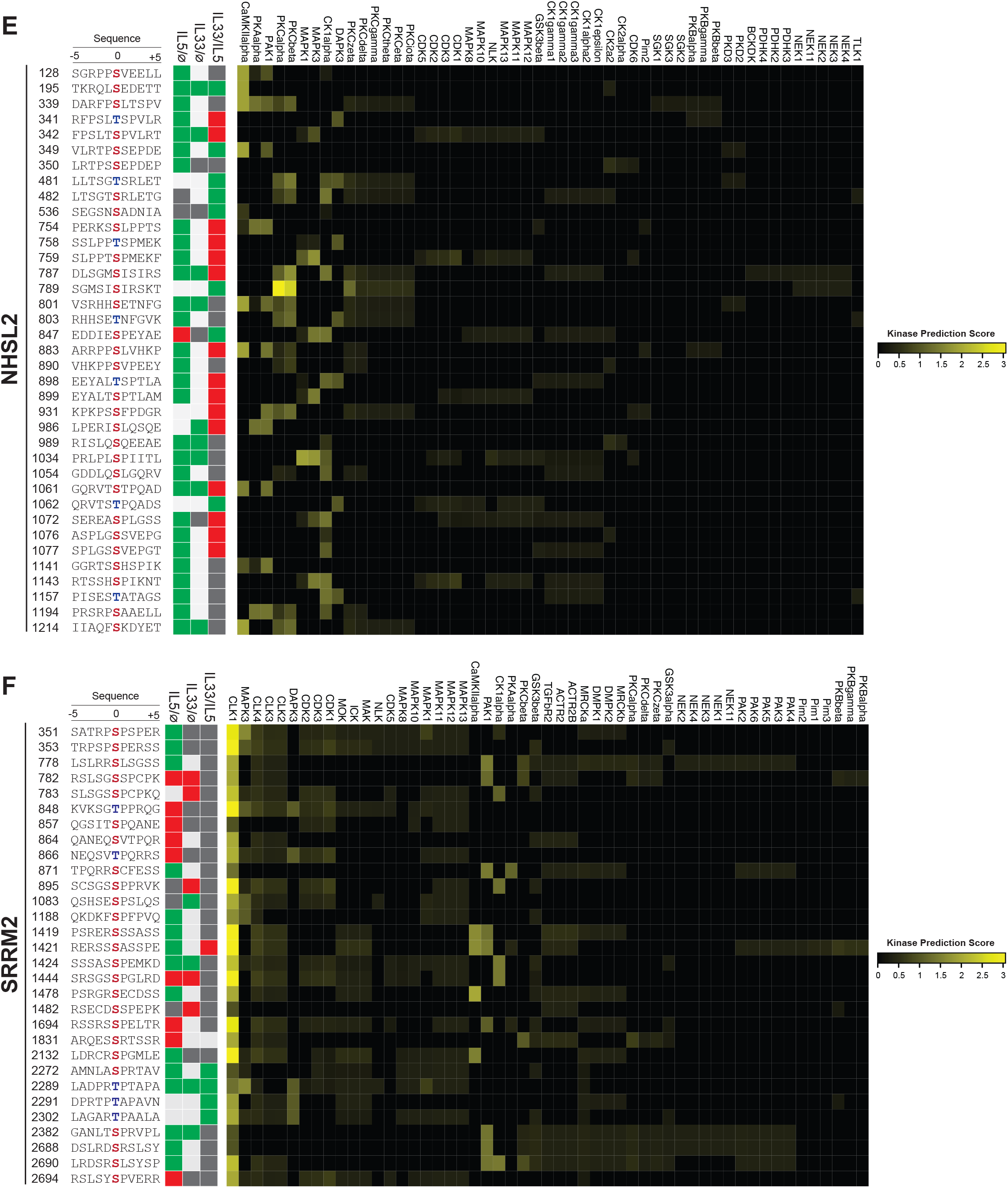
Predicted kinase regulation of highly phosphorylated multisite proteins from eosinophils. **(A-F)** Heatmaps of predicted kinase regulation of significantly changing phosphosites. The NetPhorest algorithm was used to predict kinase-substrate relationships between the indicated phosphosites. Significantly changing phosphosites for each indicated protein are listed on the left. Sequence motifs are shown on the left. The phosphoresidue is indicated in red or blue (serine/threonine). The regulation of each motif is shown as upregulation (green), downregulation (red), no significant change (dark grey), or not identified (light grey) by the indicated phosphoproteomic dataset. In the IL33 versus IL5 dataset, green corresponds to increased phosphorylation in IL33 whereases red indicates increased phosphorylation in IL5. Kinases are labeled on the horizontal axis (top) in order of their kinase prediction score labeled in yellow. The kinase prediction score scales are shown on the right for each protein.

**Figure 7:**
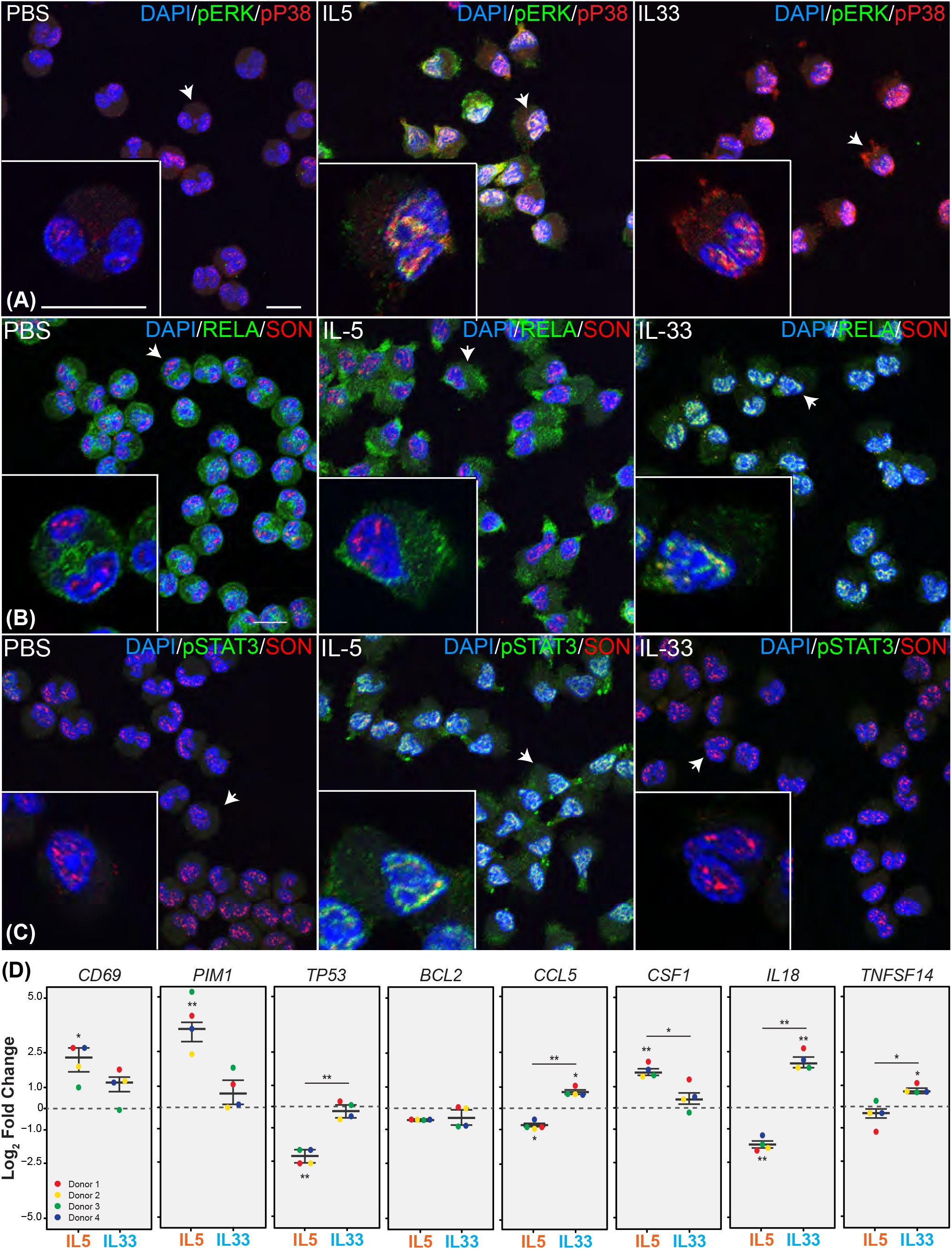
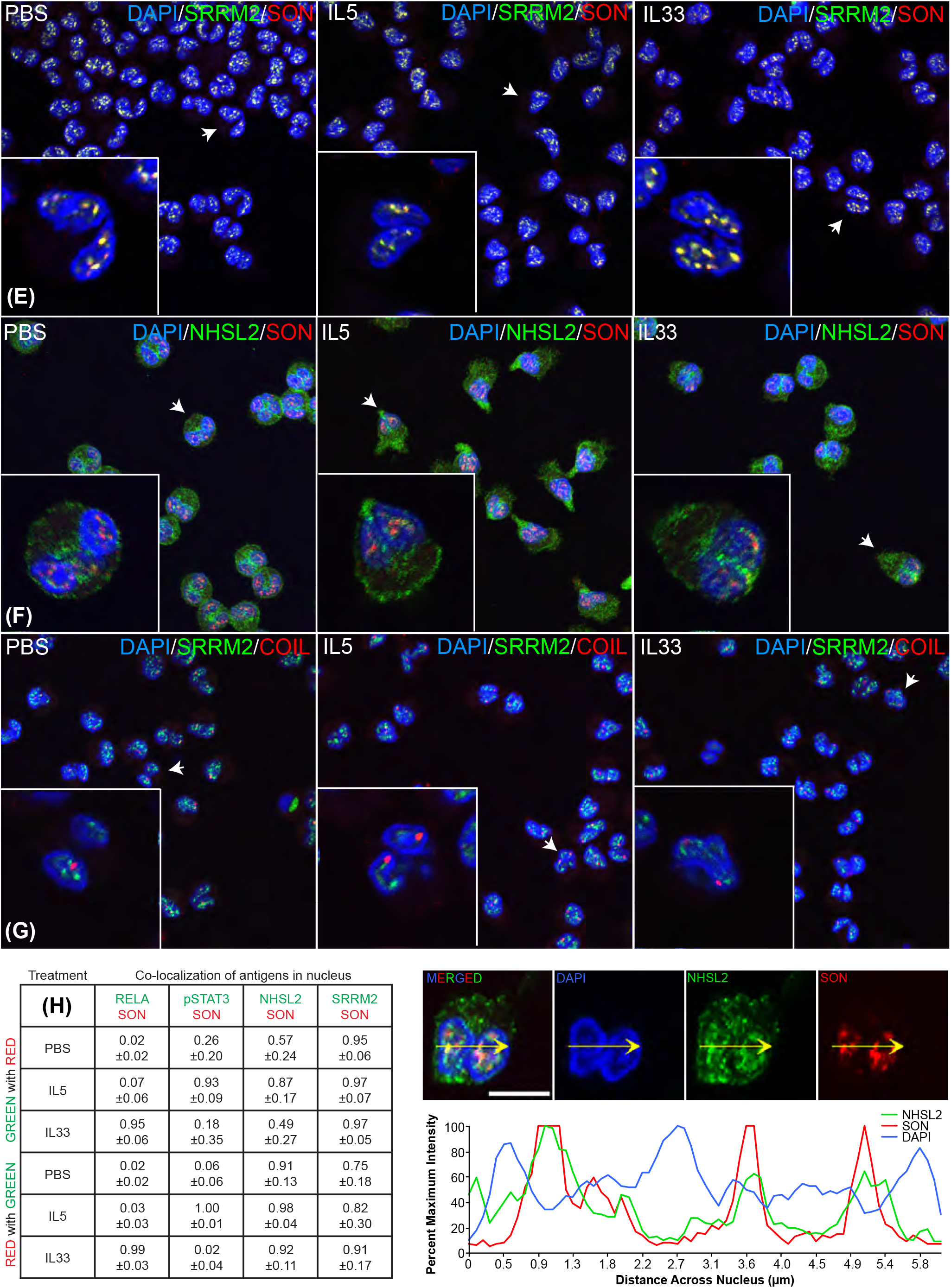
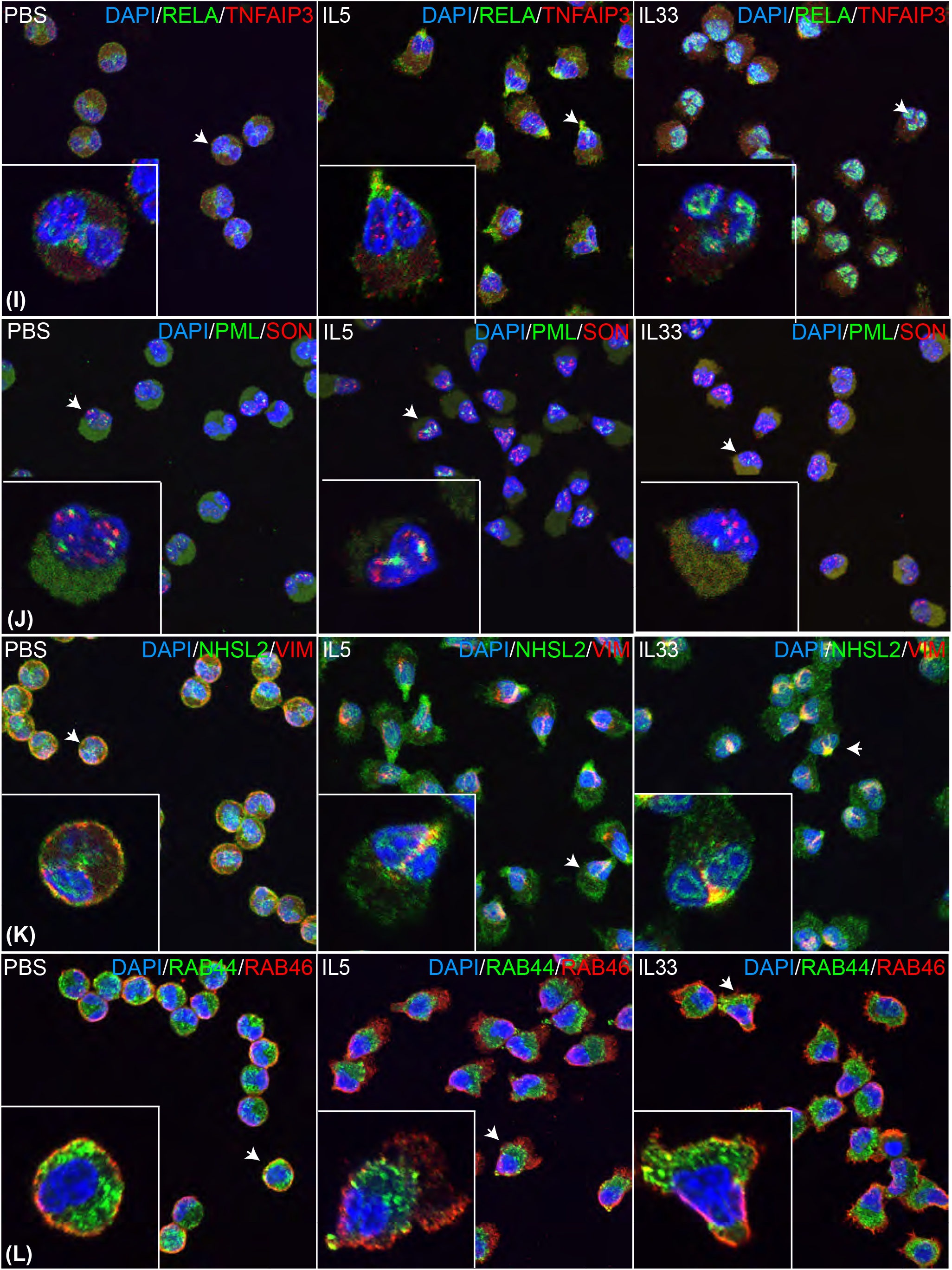
Localization of differentially phosphorylated proteins in eosinophils after 10 min of activation and effects of activation on gene expression after 2h of activation. **(A-C), (E-G)**, and **(I-L)** are confocal double immuno-fluorescence micrographs of purified eosinophils incubated for 10 ((**A-C) and (E-G, I-K)**) or 40 (**L**) min without cytokine (PBS), 50 ng/mL IL5 (IL5), or 50 ng/ml IL33 (IL33). The targets for the primary antibodies are labeled according to their color in each image. Cells were fixed with 3.7% paraformaldehyde for 10 min, quenched with 0.1 M glycine for 10 min, re-suspended in PBS, and cytospun onto poly-l-lysine-coated glass coverslips. Eosinophils were permeabilized using 0.5% SDS in PBS for 15 min. After removal of SDS, coverslips were washed, incubated with 10% bovine serum albumin (BSA) for 1 h, and incubated for 12-15 h at 4°C in primary antibodies diluted in 2% BSA and 0.1% SDS in PBS. Coverslips were washed in PBS and incubated for 1 h at room temperature with Alexa Fluor labeled secondary antibodies in 2% BSA and 0.1% SDS in PBS. Coverslips were then washed with PBS, incubated 5 min After incubation with DAPI, coverslips were then mounted on slides in Prolong Diamond anti-fade medium (InVitrogen). Sequential 0.3-μm z-step images were acquired using a confocal microscope (Nikon A1R-Si+ Confocal, 60X oil objective NA1.4) at excitation/emission wavelengths of 405/450 nm, 488/525 nm, and 555/640 nm. Laser power settings and conversion gains were kept constant for activated and non-activated cells and for control IgG staining. Images were captured with a Nikon A1 + camera, reconstructed with NIS Elements AR v4.6, and are displayed as both single slices and maximum projections (signal from all slices). In (**A-C), (E-G),** and (**I-L)**, each panel consists of a maximum projection image of 9-16 0.3-μm slices at lower magnification and an insert with a 0.3-μm slice of interest at higher magnification of the cell indicated by the arrowhead (scale bars 10 μm). **(A)** Localization of rabbit anti-pERK versus mouse anti-pP38. **(B)** Localization of rabbit anti-RELA versus mouse anti-SON. (**C)** Localization of rabbit anti-pSTAT3 versus mouse anti-SON. **(D)** Quantitative PCR of eight transcripts in eosinophils treated for 2 h with IL5 or IL33, 50 ng/ml, or no treatment. In brief, RNA from 8-10 million activated eosinophils from four separate donors was purified and RTqPCR using 25ng/μL of total RNA was performed as previously described using gene specific primers listed in **Supplementary Table 3**^31^. Readouts were normalized for amount of GAPDH and expressed as log2 change in relation to control. Results from individual donors are colored the same, and mean values are shown as a dash. Significant differences from no treatment or between the two cytokines are signified by asterisks; *, P<0.05; **, P<0.01. (**E)** Localization of rabbit anti-SRRM2 versus mouse anti-SON. (**F)** Localization of rabbit anti-NHSL2 versus mouse anti-SON. **(G)** Localization of rabbit anti-SRRM2 versus mouse anti-COIL. **(H)** Overlap of SON with other nuclear proteins. **Left**: Table of overlaps. **Right above**: Individual fluorescent images of a single z-plane of IL5-stimulated eosinophils stained for NHSL2, SON, DNA, or all three (scale bar 5 μm). **Right below**: Histograms of intensities of staining for NHSL2, SON, or DNA along the transit shown in the images above. **(I)** Localization of rabbit anti-RELA versus mouse anti-TNFAIP3. **(J)** Localization of rabbit anti-PML versus mouse anti-SON. **(K)** Localization of rabbit anti-NHSL2 versus mouse anti-VIM. **(L)** Localization of rabbit anti-RAB44 versus mouse anti-RAB46. Co-localization of mouse and rabbit antibodies in **(H)** was quantified using the General Analysis module of NIS Elements AR v4.6 software. Intensity thresholds were selected to define binary masks for the single blue (405), green (488) and red (555) channels that were applied to each 0.3 μm slice of a confocal Z-series (18-41 eosinophils in the image plane) and the masked area (μm^2^) was measured. Binary masks of the combined area of signal overlap from the green and red single channel masks were created using the expressions: “488 having 555” (combined GR) and “555 having 488” (combined RG). To limit analyses to the nucleus, additional binary masks were created using the expressions: “405 having combined GR” (combined BGR), “405 having combined RG” (combined BRG), “405 having 488” (combined BG) and “405 having 555” (combined RG). The fraction of signal overlap was determined using the following quotients: overlapping mask areas (combined BGR.Area)/(combined BG.Area) and (combined BRG.Area)/(combined BR.Area). The table in **(H)** shows the percent overlap reported as an average from 9-15 slices with the standard deviation representing variation among the slices. The adjacent pictures are confocal images of a single 0.3 μm z-slice of a cell activated with IL5 and triple stained with DAPI (blue), rabbit anti-NHSL2 (green) and mouse anti-SON (red). The merged image and individual channels are shown (scale bar, 5 μm). NIS Elements software was used to generate a line scan showing the blue, green, and red signal intensity from left to right along the yellow arrow was used to generate a plot of the % maximum signal intensity versus distance along the arrow.

Loss of TNFAIP3 in humans causes Behcet-like 1 familial autoinflammatory syndrome^66^, and its phosphorylation is known to be important for TNFAIP3 cleavage of linear polyubiquitin^67^. The enrichment of phosphorylation of potential IKKa/b kinase sites (S220, S592, S698, S729) accords with the roles of IKKs mediating NFκB-driven ubiquitin remodeling (**Figure 6A and Figure 5-XIII**). PML is a tumor suppressor that is the primary component to nuclear PML bodies, and its function is highly regulated by post-translational modifications^37^. The majority of phosphorylation changes occurred during IL5 activation, and three of these sites are strongly enriched for ATM and ATR kinase motifs (**Figure 6B**) ^68–71^. Two of the increased phosphosites (S480 and T482) are present within its disordered nuclear localization signal and are five amino acids upstream of lysine 487, that when acetylated negatively regulates the proapoptotic function of PML^72^.

The two “large RABs” of eosinophils, RAB44 and RAB46, contain long N-terminal extensions that allow direct engagement of targeted vesicles with dynein^73^. Labile RAB46 phosphosites are in the proline-rich linker between the N-terminal EF-hand and coiled-coil domains that bind dynein to the RAB domain and strongly enriched for the calciumdependent kinase CMKII motifs (**Figure 6C**). The sequence for RAB44 is not available for analysis by NetPhorest. When mapped to domains of RAB44, most labile sites are also in the proline-rich linker, which is considerably longer in RAB44 than in RAB46. Effects of RAB46 on dynein motility are controlled by calcium ion^73^. The possibility exists that calcium controls function of RAB44 and RAB46 both by binding to the N-terminal EF hands and by activation of CMKII.

Vimentin (VIM), which forms intermediate filaments, is important for nuclear stability during cell migration^74^. Three residues, S420, S430, and T436, were preferentially phosphorylated in IL33 eosinophils and are not matched by NetPhorest to any predicted kinases (**Figure 6D**). These sites are present just downstream of the intermediate filament (IF) rod domain and may be important for protein-protein dimerization^75^. Thus, information about the responsible kinases and functions of the phosphorylation events may lead to better understandings of VIM per se and of specific effects of IL33 on eosinophils.

Little is known the function of NHSL2, a member of the Nance-Horan Syndrome (NHS) protein family thought to contribute to cytoskeletal rearrangement^76,77^. Labile NHSL2 phosphosites are enriched in three cytosolic kinases, CMKIIA, PKA, and PKC (**Figure 6E**). However, all NHSL2 phosphosites show a large diversity in prediction scores, with many of the identified phosphosites not having any notable kinase motif enrichment (i.e., S350, S536, S989). The function of NHSL2 and the significance of these phosphorylations remain to be investigated.

Nuclear speckle and pre-mRNA splicing factor protein SRRM2 has been found to harbor hundreds of phosphorylation sites^78^. Upon IL5 and IL33 activation, most changing phosphosites are strongly enriched in CLK1 sites and to a lesser extent in MAPK3 and CAMKII (**Figure 6F**). These sites are present in disordered arginine-serine-rich domains. Nearly the entire protein is predicted to be highly disordered and recently these intrinsically disordered regions have been shown to be required for forming nuclear speckles^79^. Previous work on SRRM2 has found that depletion or truncation of SRRM2 dysregulates splicing through the loss of nuclear speckles leading to changes in immune cell cytokine secretion and innate immune function^79,80^. The modulation of SRRM2 phosphorylation remains unclear but may have important implications on pre-mRNA splicing outcomes during eosinophil activation.

Overall, the multisite proteins do not show a strong overlap in shared kinases. Rather, predicted kinase regulation follows the functional localization. SRRM2 and PML are enriched in sites for kinases that function in the nucleus (CLK1, ATM, ATR). CLK1, a nuclear active kinase, is regulated by AKT2 and plays an active role in spliceosome formation^81,82^. ATM and ATR are members of the PI3KK family, which also includes mTOR, and are known to regulate DNA metabolism and transcription^83^. In contrast, sites for CAMKII as enriched in the cytosolic proteins RAB46, VIM, NHSL2, and TNFAIP3. An interesting possibility is that rapid calcium signaling may be one mechanism that eosinophils have exploited to activate multiple cell-wide processes at once^84^.

### Cytokine specific relocalization of transcriptional activators and changes in gene expression

To confirm the presence and nuclear translocations of activated MAPKs, RELA, and pSTAT3 depicted in **Figure 5**, we imaged non-activated and activated eosinophils. We observed specific localization of these transcription factors in a cytokine-specific manner: non-activated eosinophils had no detectable pERK and some nuclear pP38 (**Figure 7A**). Consisted with the phosphoproteomics, nuclear pERK increased after IL5 stimulation, and nuclear pP38 increased after IL33 stimulation. Based on quantitative proteomics of eosinophils^64^, anti-pERK likely recognizes activated MAPK1 and MAPK3 and anti-pP38 mainly recognizes activated MAPK14. RELA localized to the nucleopod tip of IL5-activated eosinophils and nuclei of IL33-activated eosinophils (**Figure 7B**) whereas pSTAT3 localized to the nucleopod tip and the nucleus of IL5-activated eosinophils but was not detectable in non-treated or IL33-treated eosinophils (**Figure 7C**).

To detect possible differential changes in transcript abundance stemming from the two cytokines, we activated eosinophils with IL5 or IL33 for 2h, pelleted cells, and isolated RNA. Relative gene expression changes after two hours of IL5 or IL33 treatment were compared to non-activated eosinophils (**Figure 7D**). Differential responses were found for the four cytokines or chemokines most abundant in non-stimulated eosinophils^64^, with IL5 favoring CD69, PIM1, and CSF1 transcripts and IL33 favoring CCL5 (RANTES), IL18, and TNFSF14 (LIGHT) transcripts^85–87^. CD69 abundance increases upon activation with multiple cytokines including IL5 (and IL33) and is a hallmark of eosinophil activation^85,88,89^. PIM1 transcripts significantly increased with IL5 exposure, as has been reported previously^88,90^, whereas no significant change is measured after treatment with IL33. The significant changes are reflected in known transcriptional regulation: CCL5 and IL18 are known to be regulated through NFκB signaling^91,92^, and PIM1 and CSF1 are regulated by JAK/STAT signaling^93,94^.

### Insights into cellular architecture during activation revealed by imaging of multisite-phosphoproteins

In **Figure 7B** (IL33) and **7C** (IL5), RELA and pSTAT3 localized to dots in each nuclear lobe just inside the dense rim of DAPI-staining DNA; the dots overlapped with staining of SON, a nuclear speckle protein^95,96^. As a positive control for this overlap, we stained for SRRM2, which is known to associate with SON in nuclear speckles^95,96^ and found similar overlapping staining (**Figure 7E**). NHSL2, which has not been localized before in any cell type, was in the cytoplasm and nucleus under all three conditions (**Figure 7F**). In IL5-activated eosinophils, cytoplasmic NHSL2 was enriched in the nucleopod tip. Upon activation with either cytokine, nuclear NHSL2 had an inner rim-like distribution that co-localized with SON (**Figure 7F**). As expected, we did not find SRRM2 co-localization with COIL, a major constituent of Cajal bodies^97^ (**Figure 7G**). Interestingly, in all eosinophils in which we could clearly distinguish two nuclear lobes, each lobe had only one COIL-staining body. This is in contrast to many human cell types that contain multiple Cajal bodies. Percentages of overlap of RELA, pSTAT3, SRRM2 or NHSL2 with SON in the nucleus are given in **Figure 7H** along with histograms of signal due to DNA, NHSL2, and SON across a transit placed to cross adjacent nuclear lobes of a single eosinophil. The co-localizing proteins are found interior to rims of dense DNA as shown by the peaks in DAPI staining as the transit crosses the outer rim of the first lobe, the adjacent outer rims of the first and second lobes, and the outer rim of the second lobe.

Although IL5 did not cause RELA to move into the nucleus, RELA did translocate to the nucleopod tip (**Figure 7B and 7I**). In contrast, TNFAIP3 was in dots in the cytoplasm and nucleus of non-activated eosinophils with no obvious relocalization upon activation with either (**Figure 7I**). There were multiple PML bodies in each lobe (**Figure 7J**), and this number did not change with activation despite the changes found in its phosphorylation (**Figure 5**). The relocalization of VIM was different from that of NHSL2 (**Figure 7K**). VIM condensed from the membrane of non-activated eosinophils to the space between the bi-lobed nuclei in both IL5- and IL33-activated eosinophils. We noted no obvious differences between the localization of vimentin between IL33- and IL5-activated cells, even though there were large increases in phosphorylation at S420, S430, and T436 in IL33-activated compared to IL5-activated eosinophils (**Figure 5**).

The two “large RABs” of eosinophils, RAB44 and RAB46, localized differently (**Figure 7L**). These RABs contain long N-terminal extensions that allow direct engagement of targeted vesicles with dynein^73^. RAB44 is highly abundant in eosinophils^64^ and when knocked-out in mice impairs degranulation of mast cells^19^. RAB44 localized to the region containing crystalloid granules in both non-stimulated and cytokine-stimulated cells and to the nucleopod after stimulation with IL5. In contrast, RAB46 relocalized from the peripheral membrane to the nucleopod. These results suggest that the two RABs transport different types of vesicles, with RAB44 having the function of moving contents out of crystalloid granules. The roles of phosphorylation in causing relocalization of these proteins are not known.

## DISCUSSION

Here we report an in-depth study of the comparisons in signaling pathways utilized during IL5 and IL33 activation. We found that within the first ten minutes of cytokine exposure eosinophils show dramatic changes in their phosphoproteome, without global protein changes. After IL5 or IL33 activation, greater than 25% of the total phosphosites changed. Many of the changes were shared between both activators. A direct comparison of IL5- and IL33-treated cells demonstrated that the fold-changes at many of these sites were different.

We note that many of the most dynamic phosphosite changes are occurring in proteins with multisite phosphorylation. A general positional assessment of these sites by AlphaFold predictions revealed that many of these phosphosites, of which few are well-characterized, reside in disordered regions, even in such a highly structured protein like VIM^98^. Our data highlight the fact that multi-site phosphorylated, low complexity proteins account for much of the reshaping of the eosinophil phosphoproteome and are involved in rapid phosphorylation and engagement of proteins involved in vesicle trafficking, cytoskeletal rearrangement, splicing, and nuclear architecture. Our analysis suggests these proteins may play a more significant role in eosinophil activation than previously appreciated, and additional work is needed in defining their significance.

The network mapping of the activated pathways seen in IL5 and IL33 is rich in information on the targeted differences between the phosphorylation of the same sets of substrates, but at different residues. Our understanding of the differences in which these phosphorylation events may function is limited as a large number of these phosphosites remain uncharacterized, despite their annotation in the PhosphositePlus database^99^. Surprisingly, we find that proteins from the same family were selectively phosphorylated upon cytokine activation. For instance, SOS1 and SOS2 were distinctly phosphorylated during IL5 or IL33 activation, respectively. We also find that many of the enriched KEGG pathways, including JAK/STAT and NFκB signaling cascades, do not exhibit the traditional activating phosphorylation events, and in some cases, we see inhibitory phosphorylation sites increasing^100,101^. Despite these findings, downstream phosphorylation events indicative of pathway activation, such as tyrosine phosphorylation amongst the STATs and phosphorylation of the MAPKAP2/3K kinases, are well represented^40,62,63^. This suggests that some of the phosphorylations observed after ten minutes could be a mix of the initial signal transduction that occurred just after activation, and phosphorylations that occur as a result of cytokine commitment to an activated state. In either case, these data provide a greater understanding of how IL5 and IL33 can trigger different signaling pathways that alter downstream processes, including cell morphology and transcriptional activation and ultimately the potential responses in tissue affecting disease activity.

We provide a molecular description of the bi-lobed eosinophil nucleus and how it is impacted cytokine activation. The nucleus appears poised to become an epicenter of biological function upon cytokine exposure, with specific transcription factor translocation depending upon activation. IL5 activation results in pERK and pSTAT3 localizing to the nucleus, whereas with IL33 pP38 (MAPK14) and RELA localize to the nucleus. RELA and pSTAT3 co-localize to nuclear speckles with the splicing proteins SON and SRRM2. Surprisingly, NHSL2 also co-localizes to nuclear speckles. Remarkably, each lobe had the same appearance and presumably the same potential to act transcriptionally independent from the other with the presence of a Cajal body in each lobe, which allows for the modification of transcripts and shaping of the chromatin interaction landscape^102^. Upon nuclear localization these proteins occupy the “void volume” of the eosinophil nucleus, presumably the euchromatin that lies between the brightly stained nuclear envelope-associated heterochromatin. We speculate that the resting eosinophil nucleus remains in a poised state that can rapidly respond to extracellular stimuli.

As described in the introduction, there has been considerable success in inhibiting various cytokines and cytokine receptors to control asthma. Our detailed description of the signal transduction pathways utilized during IL5 and IL33 activation can be used to identify and develop novel therapeutics. This approach could also be used to investigate the activation states and phenotypic variation of other granulocytes. Furthermore, based on the broad coverage presented here, we expect that a global MS approach employing TMT and more targeted MS approaches can be used to target and test therapeutics that inhibit multiple cytokine pathways.

## Acknowledgements

The authors would like to thank Paul Fichtinger for purification of eosinophils from donor blood. Microscopy was performed at the University of Wisconsin-Madison Biochemistry Optical Core, which was established with support from the University of Wisconsin-Madison Department of Biochemistry Endowment.

## Author contributions

Writing: JMM, JWM, LKM, MWJ, FJF, DFM. Antibody production and characterization: JMM, DSA. Eosinophil purification, SKM. Isobaric labeling and LC-MS/MS: LKM, ASH. Microscopy: FJF. Quantitative RT-PCR, JWM. Funding and overall responsibility, JJC, DFM.

## References

1. Mack, E.A., and Pear, W.S. (2020). Transcription factor and cytokine regulation of eosinophil lineage commitment. Curr. Opin. Hematol. 27, 27–33. 10.1097/moh.0000000000000552.

2. Chusid, M.J. (2018). Eosinophils: Friends or Foes? J. Allergy Clin. Immunol. Pract. 6, 1439–1444. 10.1016/j.jaip.2018.04.031.

3. Rosenberg, H.F., Dyer, K.D., and Foster, P.S. (2013). Eosinophils: changing perspectives in health and disease. Nat. Rev. Immunol. 13, 9–22. 10.1038/nri3341.

4. Bochner, B.S. (2018). The eosinophil: For better or worse, in sickness and in health. Ann. Allergy Asthma Immunol. Off. Publ. Am. Coll. Allergy Asthma Immunol. 121, 150–155. 10.1016/j.anai.2018.02.031.

5. Wechsler, M.E., Munitz, A., Ackerman, S.J., Drake, M.G., Jackson, D.J., Wardlaw, A.J., Dougan, S.K., Berdnikovs, S., Schleich, F., Matucci, A., et al. (2021). Eosinophils in Health and Disease: A State-of-the-Art Review. Mayo Clin. Proc. 96, 2694–2707. 10.1016/j.mayocp.2021.04.025.

6. Davoine, F., and Lacy, P. (2014). Eosinophil Cytokines, Chemokines, and Growth Factors: Emerging Roles in Immunity. Front. Immunol. 5. 10.3389/fimmu.2014.00570.

7. Han, S.-T., and Mosher, D.F. (2013). IL-5 Induces Suspended Eosinophils to Undergo Unique Global Reorganization Associated with Priming. Am. J. Respir. Cell Mol. Biol. 50, 654–664. 10.1165/rcmb.2013-0181OC.

8. Bahaie, N.S., Hosseinkhani, M.R., Ge, X.N., Kang, B.N., Ha, S.G., Blumenthal, M.S., Jessberger, R., Rao, S.P., and Sriramarao, P. (2012). Regulation of Eosinophil Trafficking by SWAP-70 and Its Role in Allergic Airway Inflammation. J. Immunol. 188, 1479–1490. 10.4049/jimmunol.1102253.

9. Fettrelet, T., Gigon, L., Karaulov, A., Yousefi, S., and Simon, H.-U. (2021). The Enigma of Eosinophil Degranulation. Int. J. Mol. Sci. 22, 7091. 10.3390/ijms22137091.

10. Weller, P.F., and Spencer, L.A. (2017). Functions of tissue-resident eosinophils. Nat. Rev. Immunol. 17, 746–760. 10.1038/nri.2017.95.

11. Khoury, P., Akuthota, P., Ackerman, S.J., Arron, J.R., Bochner, B.S., Collins, M.H., Kahn, J.-E., Fulkerson, P.C., Gleich, G.J., Gopal-Srivastava, R., et al. (2018). Revisiting the NIH Taskforce on the Research needs of Eosinophil-Associated Diseases (RETREAD). J. Leukoc. Biol. 104, 69–83. 10.1002/JLB.5MR0118-028R.

12. Angulo, E.L., McKernan, E.M., Fichtinger, P.S., and Mathur, S.K. (2019). Comparison of IL-33 and IL-5 family mediated activation of human eosinophils. PLOS ONE 14, e0217807. 10.1371/journal.pone.0217807.

13. Cherry, W.B., Yoon, J., Bartemes, K.R., Iijima, K., and Kita, H. (2008). A novel IL-1 family cytokine, IL-33, potently activates human eosinophils. J. Allergy Clin. Immunol. 121, 1484–1490. 10.1016/j.jaci.2008.04.005.

14. Suzukawa, M., Koketsu, R., Iikura, M., Nakae, S., Matsumoto, K., Nagase, H., Saito, H., Matsushima, K., Ohta, K., Yamamoto, K., et al. (2008). Interleukin-33 enhances adhesion, CD11b expression and survival in human eosinophils. Lab. Invest. 88, 1245–1253. 10.1038/labinvest.2008.82.

15. Kouro, T., and Takatsu, K. (2009). IL-5- and eosinophil-mediated inflammation: from discovery to therapy. Int. Immunol. 21, 1303–1309. 10.1093/intimm/dxp102.

16. Adachi, T., and Alam, R. (1998). The mechanism of IL-5 signal transduction. Am. J. Physiol.-Cell Physiol. 275, C623–C633. 10.1152/ajpcell.1998.275.3.C623.

17. Pelaia, C., Paoletti, G., Puggioni, F., Racca, F., Pelaia, G., Canonica, G.W., and Heffler, E. (2019). Interleukin-5 in the Pathophysiology of Severe Asthma. Front. Physiol. 10, 1514. 10.3389/fphys.2019.01514.

18. Cayrol, C., and Girard, J.-P. (2014). IL-33: an alarmin cytokine with crucial roles in innate immunity, inflammation and allergy. Curr. Opin. Immunol. 31, 31–37. 10.1016/j.coi.2014.09.004.

19. Drake, L.Y., and Kita, H. (2017). IL-33: biological properties, functions, and roles in airway disease. Immunol. Rev. 278, 173–184. 10.1111/imr.12552.

20. Bouffi, C., Rochman, M., Zust, C.B., Stucke, E.M., Kartashov, A., Fulkerson, P.C., Barski, A., and Rothenberg, M.E. (2013). IL-33 Markedly Activates Murine Eosinophils by an NF-κB–Dependent Mechanism Differentially Dependent upon an IL-4–Driven Autoinflammatory Loop. J. Immunol. 191, 4317–4325. 10.4049/jimmunol.1301465.

21. Wechsler, M.E., Ruddy, M.K., Pavord, I.D., Israel, E., Rabe, K.F., Ford, L.B., Maspero, J.F., Abdulai, R.M., Hu, C.-C., Martincova, R., et al. (2021). Efficacy and Safety of Itepekimab in Patients with Moderate-to-Severe Asthma. N. Engl. J. Med. 385, 1656–1668. 10.1056/NEJMoa2024257.

22. Haldar, P., Brightling, C.E., Hargadon, B., Gupta, S., Monteiro, W., Sousa, A., Marshall, R.P., Bradding, P., Green, R.H., Wardlaw, A.J., et al. (2009). Mepolizumab and exacerbations of refractory eosinophilic asthma. N. Engl. J. Med. 360, 973–984. 10.1056/NEJMoa0808991.

23. Kelly, E.A., Esnault, S., Liu, L.Y., Evans, M.D., Johansson, M.W., Mathur, S., Mosher, D.F., Denlinger, L.C., and Jarjour, N.N. (2017). Mepolizumab Attenuates Airway Eosinophil Numbers, but Not Their Functional Phenotype, in Asthma. Am. J. Respir. Crit. Care Med. 196, 1385–1395. 10.1164/rccm.201611-2234OC.

24. Nair, P., Pizzichini, M.M.M., Kjarsgaard, M., Inman, M.D., Efthimiadis, A., Pizzichini, E., Hargreave, F.E., and O’Byrne, P.M. (2009). Mepolizumab for prednisone-dependent asthma with sputum eosinophilia. N. Engl. J. Med. 360, 985–993. 10.1056/NEJMoa0805435.

25. Senko, M.W., Remes, P.M., Canterbury, J.D., Mathur, R., Song, Q., Eliuk, S.M., Mullen, C., Earley, L., Hardman, M., Blethrow, J.D., et al. (2013). Novel parallelized quadrupole/linear ion trap/Orbitrap tribrid mass spectrometer improving proteome coverage and peptide identification rates. Anal. Chem. 85, 11710–11714. 10.1021/ac403115c.

26. Riley, N.M., and Coon, J.J. (2016). Phosphoproteomics in the Age of Rapid and Deep Proteome Profiling. Anal. Chem. 88, 74–94. 10.1021/acs.analchem.5b04123.

27. Shishkova, E., Hebert, A.S., and Coon, J.J. (2016). Now, More Than Ever, Proteomics Needs Better Chromatography. Cell Syst. 3, 321–324. 10.1016/j.cels.2016.10.007.

28. Shishkova, E., Hebert, A.S., Westphall, M.S., and Coon, J.J. (2018). Ultra-High Pressure (>30,000 psi) Packing of Capillary Columns Enhancing Depth of Shotgun Proteomic Analyses. Anal. Chem. 90, 11503–11508. 10.1021/acs.analchem.8b02766.

29. Muehlbauer, L.K., Wei, T., Shishkova, E., Coon, J.J., and Lambert, P.F. (2022). IQGAP1 and RNA Splicing in the Context of Head and Neck via Phosphoproteomics. J. Proteome Res. 21, 2211–2223. 10.1021/acs.jproteome.2c00309.

30. Bortnov, V., Annis, D.S., Fogerty, F.J., Barretto, K.T., Turton, K.B., and Mosher, D.F. (2018). Myeloid-derived growth factor is a resident endoplasmic reticulum protein. J. Biol. Chem. 293, 13166–13175. 10.1074/jbc.AC118.002052.

31. Mabin, J.W., Lewis, P.W., Brow, D.A., and Dvinge, H. (2021). Human spliceosomal snRNA sequence variants generate variant spliceosomes. RNA 27, 1186–1203. 10.1261/rna.078768.121.

32. Rieckmann, J.C., Geiger, R., Hornburg, D., Wolf, T., Kveler, K., Jarrossay, D., Sallusto, F., Shen-Orr, S.S., Lanzavecchia, A., Mann, M., et al. (2017). Social network architecture of human immune cells unveiled by quantitative proteomics. Nat. Immunol. 18, 583–593. 10.1038/ni.3693.

33. el Benna, J., Faust, L.P., and Babior, B.M. (1994). The phosphorylation of the respiratory burst oxidase component p47phox during neutrophil activation. Phosphorylation of sites recognized by protein kinase C and by proline-directed kinases. J. Biol. Chem. 269, 23431–23436.

34. Rauniyar, N., and Yates, J.R.I. (2014). Isobaric Labeling-Based Relative Quantification in Shotgun Proteomics. J. Proteome Res. 13, 5293–5309. 10.1021/pr500880b.

35. Kanehisa, M. (2002). The KEGG Database. In Novartis Foundation Symposia, G. Bock and J. A. Goode, eds. (John Wiley & Sons, Ltd), pp. 91–103. 10.1002/0470857897.ch8.

36. Kanehisa, M., Furumichi, M., Tanabe, M., Sato, Y., and Morishima, K. (2017). KEGG: new perspectives on genomes, pathways, diseases and drugs. Nucleic Acids Res. 45, D353–D361. 10.1093/nar/gkw1092.

37. Martens, M., Ammar, A., Riutta, A., Waagmeester, A., Slenter, D.N., Hanspers, K., A. Miller, R., Digles, D., Lopes, E.N., Ehrhart, F., et al. (2020). WikiPathways: connecting communities. Nucleic Acids Res. 49, D613–D621. 10.1093/nar/gkaa1024.

38. Pinto, S.M., Subbannayya, Y., Rex, D.A.B., Raju, R., Chatterjee, O., Advani, J., Radhakrishnan, A., Keshava Prasad, T.S., Wani, M.R., and Pandey, A. (2018). A network map of IL-33 signaling pathway. J. Cell Commun. Signal. 12, 615–624. 10.1007/s12079-018-0464-4.

39. Rex, D.A.B., Subbannayya, Y., Modi, P.K., Palollathil, A., Gopalakrishnan, L., Bhandary, Y.P., Prasad, T.S.K., and Pinto, S.M. (2022). Temporal Quantitative Phosphoproteomics Profiling of Interleukin-33 Signaling Network Reveals Unique Modulators of Monocyte Activation. Cells 11, 138. 10.3390/cells11010138.

40. Turton, K.B., Annis, D.S., Rui, L., Esnault, S., and Mosher, D.F. (2015). Ratios of Four STAT3 Splice Variants in Human Eosinophils and Diffuse Large B Cell Lymphoma Cells. PloS One 10, e0127243. 10.1371/journal.pone.0127243.

41. Putz, E.M., Gotthardt, D., and Sexl, V. (2014). STAT1-S727 - the license to kill. OncoImmunology 3, e955441. 10.4161/21624011.2014.955441.

42. Sadzak, I., Schiff, M., Gattermeier, I., Glinitzer, R., Sauer, I., Saalmüller, A., Yang, E., Schaljo, B., and Kovarik, P. (2008). Recruitment of Stat1 to chromatin is required for interferon-induced serine phosphorylation of Stat1 transactivation domain. Proc. Natl. Acad. Sci. 105, 8944–8949. 10.1073/pnas.0801794105.

43. Medunjanin, S., Schleithoff, L., Fiegehenn, C., Weinert, S., Zuschratter, W., and Braun-Dullaeus, R.C. (2016). GSK-3β controls NF-kappaB activity via IKKγ/NEMO. Sci. Rep. 6, 38553. 10.1038/srep38553.

44. Cartwright, T., Perkins, N.D., and L. Wilson, C. (2016). NFKB1: a suppressor of inflammation, ageing and cancer. FEBS J. 283, 1812–1822. 10.1111/febs.13627.

45. Demarchi, F., Bertoli, C., Sandy, P., and Schneider, C. (2003). Glycogen synthase kinase-3 beta regulates NF-kappa B1/p105 stability. J. Biol. Chem. 278, 39583–39590. 10.1074/jbc.M305676200.

46. Xiao, G., Fong, A., and Sun, S.-C. (2004). Induction of p100 Processing by NF-κB- inducing Kinase Involves Docking IκB Kinase α (IKKα) to p100 and IKKα-mediated Phosphorylation *. J. Biol. Chem. 279, 30099–30105. 10.1074/jbc.M401428200.

47. Qu, Z., Qing, G., Rabson, A., and Xiao, G. (2004). Tax Deregulation of NF-κB2 p100 Processing Involves Both β-TrCP-dependent and -independent Mechanisms *. J. Biol. Chem. 279, 44563–44572. 10.1074/jbc.M403689200.

48. Wick, M.J., Dong, L.Q., Riojas, R.A., Ramos, F.J., and Liu, F. (2000). Mechanism of phosphorylation of protein kinase B/Akt by a constitutively active 3-phosphoinositide-dependent protein kinase-1. J. Biol. Chem. 275, 40400–40406. 10.1074/jbc.M003937200.

49. Bellacosa, A., Chan, T.O., Ahmed, N.N., Datta, K., Malstrom, S., Stokoe, D., McCormick, F., Feng, J., and Tsichlis, P. (1998). Akt activation by growth factors is a multiple-step process: the role of the PH domain. Oncogene 17, 313–325. 10.1038/sj.onc.1201947.

50. Zhang, Y., Wang, X., Qin, X., Wang, X., Liu, F., White, E., and Zheng, X.F.S. (2015). PP2AC Level Determines Differential Programming of p38-TSC-mTOR Signaling and Therapeutic Response to p38-Targeted Therapy in Colorectal Cancer. eBioMedicine 2, 1944–1956. 10.1016/j.ebiom.2015.11.031.

51. Antonia, R.J., Castillo, J., Herring, L.E., Serafin, D.S., Liu, P., Graves, L.M., Baldwin, A.S., and Hagan, R.S. (2019). TBK1 Limits mTORC1 by Promoting Phosphorylation of Raptor Ser877. Sci. Rep. 9, 13470. 10.1038/s41598-019-49707-8.

52. Foster, K.G., Acosta-Jaquez, H.A., Romeo, Y., Ekim, B., Soliman, G.A., Carriere, A., Roux, P.P., Ballif, B.A., and Fingar, D.C. (2010). Regulation of mTOR Complex 1 (mTORC1) by Raptor Ser863 and Multisite Phosphorylation *. J. Biol. Chem. 285, 80–94. 10.1074/jbc.M109.029637.

53. Carrière, A., Cargnello, M., Julien, L.-A., Gao, H., Bonneil, É., Thibault, P., and Roux, P.P. (2008). Oncogenic MAPK Signaling Stimulates mTORC1 Activity by Promoting RSK-Mediated Raptor Phosphorylation. Curr. Biol. 18, 1269–1277. 10.1016/j.cub.2008.07.078.

54. Baljuls, A., Schmitz, W., Mueller, T., Zahedi, R.P., Sickmann, A., Hekman, M., and Rapp, U.R. (2008). Positive Regulation of A-RAF by Phosphorylation of Isoform-specific Hinge Segment and Identification of Novel Phosphorylation Sites *. J. Biol. Chem. 283, 27239–27254. 10.1074/jbc.M801782200.

55. Zheng, C.F., and Guan, K.L. (1994). Activation of MEK family kinases requires phosphorylation of two conserved Ser/Thr residues. EMBO J. 13, 1123–1131. 10.1002/j.1460-2075.1994.tb06361.x.

56. Wang, L., Xue, J., Zadorozny, E.V., and Robinson, L.J. (2008). G-CSF stimulates Jak2-dependent Gab2 phosphorylation leading to Erk1/2 activation and cell proliferation. Cell. Signal. 20, 1890–1899. 10.1016/j.cellsig.2008.06.018.

57. Margolis, B., and Skolnik, E.Y. (1994). Activation of Ras by receptor tyrosine kinases. J. Am. Soc. Nephrol. JASN 5, 1288–1299. 10.1681/ASN.V561288.

58. Saha, M., Carriere, A., Cheerathodi, M., Zhang, X., Lavoie, G., Rush, J., Roux, P.P., and Ballif, B.A. (2012). RSK phosphorylates SOS1 creating 14-3-3-docking sites and negatively regulating MAPK activation. Biochem. J. 447, 159–166. 10.1042/BJ20120938.

59. Eisenhardt, A.E., Sprenger, A., Röring, M., Herr, R., Weinberg, F., Köhler, M., Braun, S., Orth, J., Diedrich, B., Lanner, U., et al. (2016). Phospho-proteomic analyses of B-Raf protein complexes reveal new regulatory principles. Oncotarget 7, 26628–26652. 10.18632/oncotarget.8427.

60. Raingeaud, J., Gupta, S., Rogers, J.S., Dickens, M., Han, J., Ulevitch, R.J., and Davis, R.J. (1995). Pro-inflammatory Cytokines and Environmental Stress Cause p38 Mitogen-activated Protein Kinase Activation by Dual Phosphorylation on Tyrosine and Threonine (*). J. Biol. Chem. 270, 7420–7426. 10.1074/jbc.270.13.7420.

61. McLaughlin, M.M., Kumar, S., McDonnell, P.C., Horn, S.V., Lee, J.C., Livi, G.P., and Young, P.R. (1996). Identification of Mitogen-activated Protein (MAP) Kinase-activated Protein Kinase-3, a Novel Substrate of CSBP p38 MAP Kinase (*). J. Biol. Chem. 271, 8488–8492. 10.1074/jbc.271.14.8488.

62. Meng, W., Swenson, L.L., Fitzgibbon, M.J., Hayakawa, K., ter Haar, E., Behrens, A.E., Fulghum, J.R., and Lippke, J.A. (2002). Structure of Mitogen-activated Protein Kinase-activated Protein (MAPKAP) Kinase 2 Suggests a Bifunctional Switch That Couples Kinase Activation with Nuclear Export. J. Biol. Chem. 277, 37401–37405. 10.1074/jbc.C200418200.

63. Ben-Levy, R., Leighton, I.A., Doza, Y.N., Attwood, P., Morrice, N., Marshall, C.J., and Cohen, P. (1995). Identification of novel phosphorylation sites required for activation of MAPKAP kinase-2. EMBO J. 14, 5920–5930. 10.1002/j.1460-2075.1995.tb00280.x.

64. Wilkerson, E.M., Johansson, M.W., Hebert, A.S., Westphall, M.S., Mathur, S.K., Jarjour, N.N., Schwantes, E.A., Mosher, D.F., and Coon, J.J. (2016). The Peripheral Blood Eosinophil Proteome. J. Proteome Res. 15, 1524–1533. 10.1021/acs.jproteome.6b00006.

65. Horn, H., Schoof, E.M., Kim, J., Robin, X., Miller, M.L., Diella, F., Palma, A., Cesareni, G., Jensen, L.J., and Linding, R. (2014). KinomeXplorer: an integrated platform for kinome biology studies. Nat. Methods 11, 603–604. 10.1038/nmeth.2968.

66. Steiner, A., Harapas, C.R., Masters, S.L., and Davidson, S. (2018). An Update on Autoinflammatory Diseases: Relopathies. Curr. Rheumatol. Rep. 20, 39. 10.1007/s11926-018-0749-x.

67. Wertz, I.E., Newton, K., Seshasayee, D., Kusam, S., Lam, C., Zhang, J., Popovych, N., Helgason, E., Schoeffler, A., Jeet, S., et al. (2016). Erratum: Phosphorylation and linear ubiquitin direct A20 inhibition of inflammation. Nature 532, 402. 10.1038/nature16541.

68. Vereecke, L., Beyaert, R., and Loo, G. van (2009). The ubiquitin-editing enzyme A20 (TNFAIP3) is a central regulator of immunopathology. Trends Immunol. 30, 383–391. 10.1016/j.it.2009.05.007.

69. Das, T., Chen, Z., Hendriks, R.W., and Kool, M. (2018). A20/Tumor Necrosis Factor α - Induced Protein 3 in Immune Cells Controls Development of Autoinflammation and Autoimmunity: Lessons from Mouse Models. Front. Immunol. 9.

70. PML: Regulation and multifaceted function beyond tumor suppression | Cell & Bioscience | Full Text https://cellandbioscience.biomedcentral.com/articles/10.1186/s13578-018-0204-8.

71. Guan, D., and Kao, H.-Y. (2015). The function, regulation and therapeutic implications of the tumor suppressor protein, PML. Cell Biosci. 5, 60. 10.1186/s13578-015-0051-9.

72. Hayakawa, F., Abe, A., Kitabayashi, I., Pandolfi, P.P., and Naoe, T. (2008). Acetylation of PML is involved in histone deacetylase inhibitor-mediated apoptosis. J. Biol. Chem. 283, 24420–24425. 10.1074/jbc.M802217200.

73. Wang, Y., Huynh, W., Skokan, T.D., Lu, W., Weiss, A., and Vale, R.D. (2019). CRACR2a is a calcium-activated dynein adaptor protein that regulates endocytic traffic. J. Cell Biol. 218, 1619–1633. 10.1083/jcb.201806097.

74. Patteson, A.E., Vahabikashi, A., Pogoda, K., Adam, S.A., Mandal, K., Kittisopikul, M., Sivagurunathan, S., Goldman, A., Goldman, R.D., and Janmey, P.A. (2019). Vimentin protects cells against nuclear rupture and DNA damage during migration. J. Cell Biol. 10.1083/jcb.201902046.

75. Snider, N.T., Ku, N.-O., and Omary, M.B. (2018). The sweet side of vimentin. eLife 7, e35336. 10.7554/eLife.35336.

76. Turton, K.B. (2018). Case Studies of Cryptic Proteins Contributing to Shape Change in Eosinophils: Workflows to Investigate STAT3, LOCGEF and NHSL2.

77. Brooks, S.P., Coccia, M., Tang, H.R., Kanuga, N., Machesky, L.M., Bailly, M., Cheetham, M.E., and Hardcastle, A.J. (2010). The Nance–Horan syndrome protein encodes a functional WAVE homology domain (WHD) and is important for co-ordinating actin remodelling and maintaining cell morphology. Hum. Mol. Genet. 19, 2421–2432. 10.1093/hmg/ddq125.

78. Ilik, İ.A., and Aktaş, T. (2022). Nuclear speckles: dynamic hubs of gene expression regulation. FEBS J. 289, 7234–7245. 10.1111/febs.16117.

79. Xu, S., Lai, S.-K., Sim, D.Y., Ang, W.S.L., Li, H.Y., and Roca, X. (2022). SRRM2 organizes splicing condensates to regulate alternative splicing. Nucleic Acids Res. 50, 8599–8614. 10.1093/nar/gkac669.

80. Whisenant, T.C., Peralta, E.R., Aarreberg, L.D., Gao, N.J., Head, S.R., Ordoukhanian, P., Williamson, J.R., and Salomon, D.R. (2015). The Activation-Induced Assembly of an RNA/Protein Interactome Centered on the Splicing Factor U2AF2 Regulates Gene Expression in Human CD4 T Cells. PloS One 10, e0144409. 10.1371/journal.pone.0144409.

81. Jiang, K., Patel, N.A., Watson, J.E., Apostolatos, H., Kleiman, E., Hanson, O., Hagiwara, M., and Cooper, D.R. (2009). Akt2 regulation of Cdc2-like kinases (Clk/Sty), serine/arginine-rich (SR) protein phosphorylation, and insulin-induced alternative splicing of PKCbetaII messenger ribonucleic acid. Endocrinology 150, 2087–2097. 10.1210/en.2008-0818.

82. Aubol, B.E., Wu, G., Keshwani, M.M., Movassat, M., Fattet, L., Hertel, K.J., Fu, X.-D., and Adams, J.A. (2016). Release of SR Proteins from CLK1 by SRPK1: A Symbiotic Kinase System for Phosphorylation Control of Pre-mRNA Splicing. Mol. Cell 63, 218–228. 10.1016/j.molcel.2016.05.034.

83. Burger, K., Ketley, R.F., and Gullerova, M. (2019). Beyond the Trinity of ATM, ATR, and DNA-PK: Multiple Kinases Shape the DNA Damage Response in Concert With RNA Metabolism. Front. Mol. Biosci. 6, 61. 10.3389/fmolb.2019.00061.

84. Son, K., Hussain, A., Sehmi, R., and Janssen, L. (2021). The Cycling of Intracellular Calcium Released in Response to Fluid Shear Stress Is Critical for Migration-Associated Actin Reorganization in Eosinophils. Cells 10, 157. 10.3390/cells10010157.

85. Johansson, M.W. (2014). Activation states of blood eosinophils in asthma. Clin. Exp. Allergy J. Br. Soc. Allergy Clin. Immunol. 44, 482–498. 10.1111/cea.12292.

86. Larose, M.-C., Archambault, A.-S., Provost, V., Laviolette, M., and Flamand, N. (2017). Regulation of Eosinophil and Group 2 Innate Lymphoid Cell Trafficking in Asthma. Front. Med. 4, 136. 10.3389/fmed.2017.00136.

87. Manresa, M.C., Miki, H., Miller, J., Okamoto, K., Dobaczewska, K., Herro, R., Gupta, R.K., Kurten, R., Aceves, S.S., and Croft, M. (2022). A Deficiency in the Cytokine TNFSF14/LIGHT Limits Inflammation and Remodeling in Murine Eosinophilic Esophagitis. J. Immunol. Baltim. Md 1950, ji2200326. 10.4049/jimmunol.2200326.

88. Bates, M.E., Liu, L.Y., Esnault, S., Stout, B.A., Fonkem, E., Kung, V., Sedgwick, J.B., Kelly, E.A.B., Bates, D.M., Malter, J.S., et al. (2004). Expression of interleukin-5- and granulocyte macrophage-colony-stimulating factor-responsive genes in blood and airway eosinophils. Am. J. Respir. Cell Mol. Biol. 30, 736–743. 10.1165/rcmb.2003-0234OC.

89. Andreone, S., Spadaro, F., Buccione, C., Mancini, J., Tinari, A., Sestili, P., Gambardella, A.R., Lucarini, V., Ziccheddu, G., Parolini, I., et al. (2019). IL-33 Promotes CD11b/CD18-Mediated Adhesion of Eosinophils to Cancer Cells and Synapse-Polarized Degranulation Leading to Tumor Cell Killing. Cancers 11, 1664. 10.3390/cancers11111664.

90. Stout, B.A., Bates, M.E., Liu, L.Y., Farrington, N.N., and Bertics, P.J. (2004). IL-5 and granulocyte-macrophage colony-stimulating factor activate STAT3 and STAT5 and promote Pim-1 and cyclin D3 protein expression in human eosinophils. J. Immunol. Baltim. Md 1950 173, 6409–6417. 10.4049/jimmunol.173.10.6409.

91. Liu, X., Shah, A., Gangwani, M.R., Silverstein, P.S., Fu, M., and Kumar, A. (2014). HIV-1 Nef induces CCL5 production in astrocytes through p38-MAPK and PI3K/Akt pathway and utilizes NF-kB, CEBP and AP-1 transcription factors. Sci. Rep. 4, 4450. 10.1038/srep04450.

92. Suk, K., Yeou Kim, S., and Kim, H. (2001). Regulation of IL-18 production by IFN gamma and PGE2 in mouse microglial cells: involvement of NF-kB pathway in the regulatory processes. Immunol. Lett. 77, 79–85. 10.1016/s0165-2478(01)00209-7.

93. Tsuchimoto, D., Tojo, A., and Asano, S. (2004). A mechanism of transcriptional regulation of the CSF-1 gene by interferon-gamma. Immunol. Invest. 33, 397–405. 10.1081/imm-200038662.

94. Narlik-Grassow, M., Blanco-Aparicio, C., and Carnero, A. (2014). The PIM family of serine/threonine kinases in cancer. Med. Res. Rev. 34, 136–159. 10.1002/med.21284.

95. Ilik, İ.A., Malszycki, M., Lübke, A.K., Schade, C., Meierhofer, D., and Aktaş, T. (2020). SON and SRRM2 are essential for nuclear speckle formation. eLife 9, e60579. 10.7554/eLife.60579.

96. Sharma, A., Takata, H., Shibahara, K., Bubulya, A., and Bubulya, P.A. (2010). Son Is Essential for Nuclear Speckle Organization and Cell Cycle Progression. Mol. Biol. Cell 21, 650–663. 10.1091/mbc.e09-02-0126.

97. Machyna, M., Heyn, P., and Neugebauer, K.M. (2013). Cajal bodies: where form meets function. Wiley Interdiscip. Rev. RNA 4, 17–34. 10.1002/wrna.1139.

98. Jumper, J., Evans, R., Pritzel, A., Green, T., Figurnov, M., Ronneberger, O., Tunyasuvunakool, K., Bates, R., Žídek, A., Potapenko, A., et al. (2021). Highly accurate protein structure prediction with AlphaFold. Nature 596, 583–589. 10.1038/s41586-021-03819-2.

99. Hornbeck, P.V., Zhang, B., Murray, B., Kornhauser, J.M., Latham, V., and Skrzypek, E. (2015). PhosphoSitePlus, 2014: mutations, PTMs and recalibrations. Nucleic Acids Res. 43, D512–D520. 10.1093/nar/gku1267.

100. Feng, J., Witthuhn, B.A., Matsuda, T., Kohlhuber, F., Kerr, I.M., and Ihle, J.N. (1997). Activation of Jak2 catalytic activity requires phosphorylation of Y1007 in the kinase activation loop. Mol. Cell. Biol. 17, 2497–2501. 10.1128/MCB.17.5.2497.

101. Argetsinger, L.S., Kouadio, J.-L.K., Steen, H., Stensballe, A., Jensen, O.N., and Carter-Su, C. (2004). Autophosphorylation of JAK2 on tyrosines 221 and 570 regulates its activity. Mol. Cell. Biol. 24, 4955–4967. 10.1128/MCB.24.11.4955-4967.2004.

102. Sawyer, I.A., Sturgill, D., Sung, M.-H., Hager, G.L., and Dundr, M. (2016). Cajal body function in genome organization and transcriptome diversity. BioEssays News Rev. Mol. Cell. Dev. Biol. 38, 1197–1208. 10.1002/bies.201600144.

103. Metz, K.S., Deoudes, E.M., Berginski, M.E., Jimenez-Ruiz, I., Aksoy, B.A., Hammerbacher, J., Gomez, S.M., and Phanstiel, D.H. (2018). Coral: Clear and Customizable Visualization of Human Kinome Data. Cell Syst. 7, 347–350.e1. 10.1016/j.cels.2018.07.001.

104. Barretto, K.T., Swanson, C.M., Nguyen, C.L., Annis, D.S., Esnault, S.J., Mosher, D.F., and Johansson, M.W. (2018). Control of cytokine-driven eosinophil migratory behavior by TGF-beta-induced protein (TGFBI) and periostin. PLOS ONE 13, e0201320. 10.1371/journal.pone.0201320.

105. Cox, J., and Mann, M. (2008). MaxQuant enables high peptide identification rates, individualized p.p.b.-range mass accuracies and proteome-wide protein quantification. Nat. Biotechnol. 26, 1367–1372. 10.1038/nbt.1511.

106. Cox, J., Neuhauser, N., Michalski, A., Scheltema, R.A., Olsen, J.V., and Mann, M. (2011). Andromeda: A Peptide Search Engine Integrated into the MaxQuant Environment. J. Proteome Res. 10, 1794–1805. 10.1021/pr101065j.

107. Tyanova, S., Temu, T., Sinitcyn, P., Carlson, A., Hein, M.Y., Geiger, T., Mann, M., and Cox, J. (2016). The Perseus computational platform for comprehensive analysis of (prote)omics data. Nat. Methods 13, 731–740. 10.1038/nmeth.3901.

108. Maurer, L.M., Tomasini-Johansson, B.R., Ma, W., Annis, D.S., Eickstaedt, N.L., Ensenberger, M.G., Satyshur, K.A., and Mosher, D.F. (2010). Extended Binding Site on Fibronectin for the Functional Upstream Domain of Protein F1 of Streptococcus pyogenes. J. Biol. Chem. 285, 41087–41099. 10.1074/jbc.M110.153692.

109. Bortnov, V., Tonelli, M., Lee, W., Lin, Z., Annis, D.S., Demerdash, O.N., Bateman, A., Mitchell, J.C., Ge, Y., Markley, J.L., et al. (2019). Solution structure of human myeloid-derived growth factor suggests a conserved function in the endoplasmic reticulum. Nat. Commun. 10, 5612. 10.1038/s41467-019-13577-5.

110. PrimerBank: a PCR primer database for quantitative gene expression analysis, 2012 update | Nucleic Acids Research | Oxford Academic https://academic.oup.com/nar/article/40/D1/D1144/2902573.

