## Supplemental Data for "Molecular anatomy of eosinophil activation by IL5 and IL33"

**Molecular anatomy of eosinophil activation by IL5 and IL33: Supplemental Material**

**Running Title:** Molecular Anatomy of Eosinophil Activation

### Supplemental Figure 1

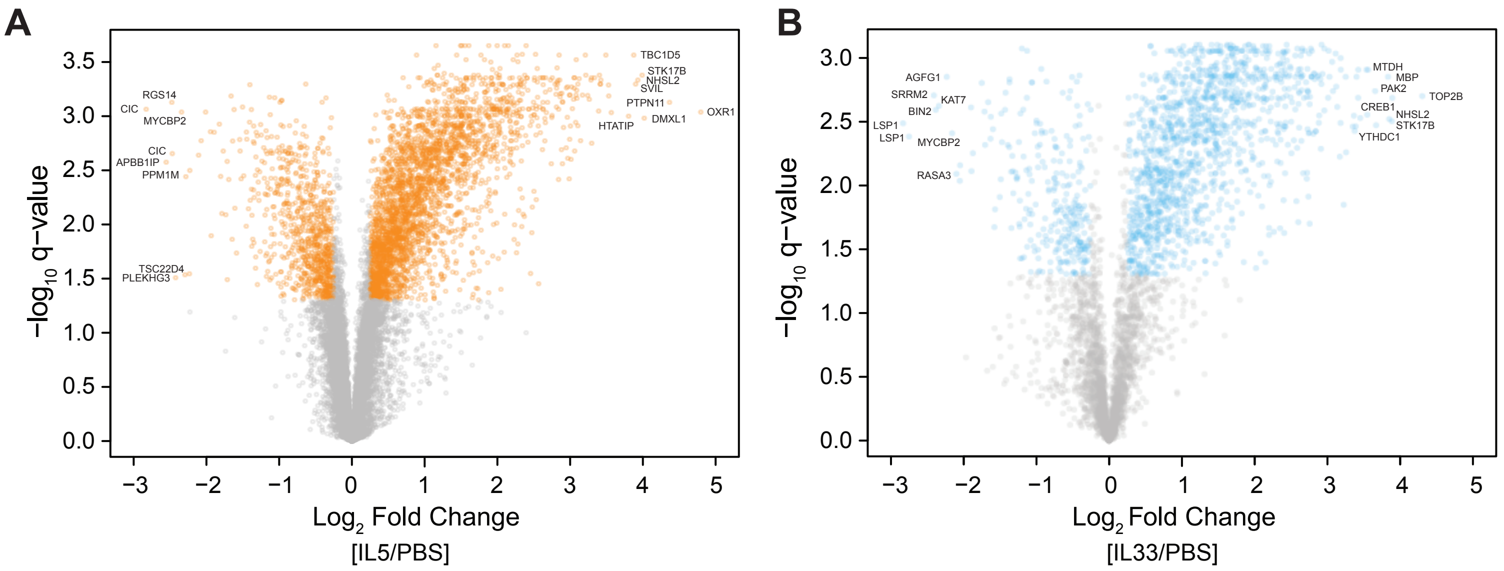


### Supplemental Figure 2

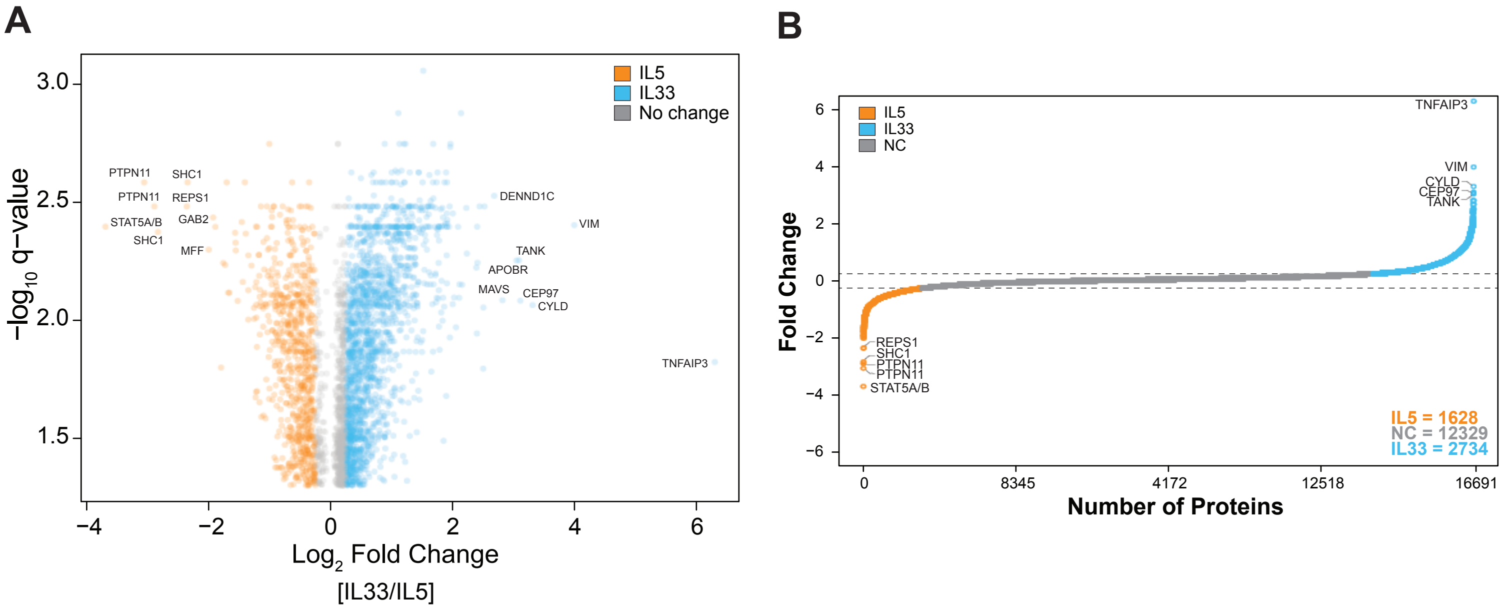

### Supplemental Figure 3

#### Mass Spectrometry Identified Eosinophil Kinases

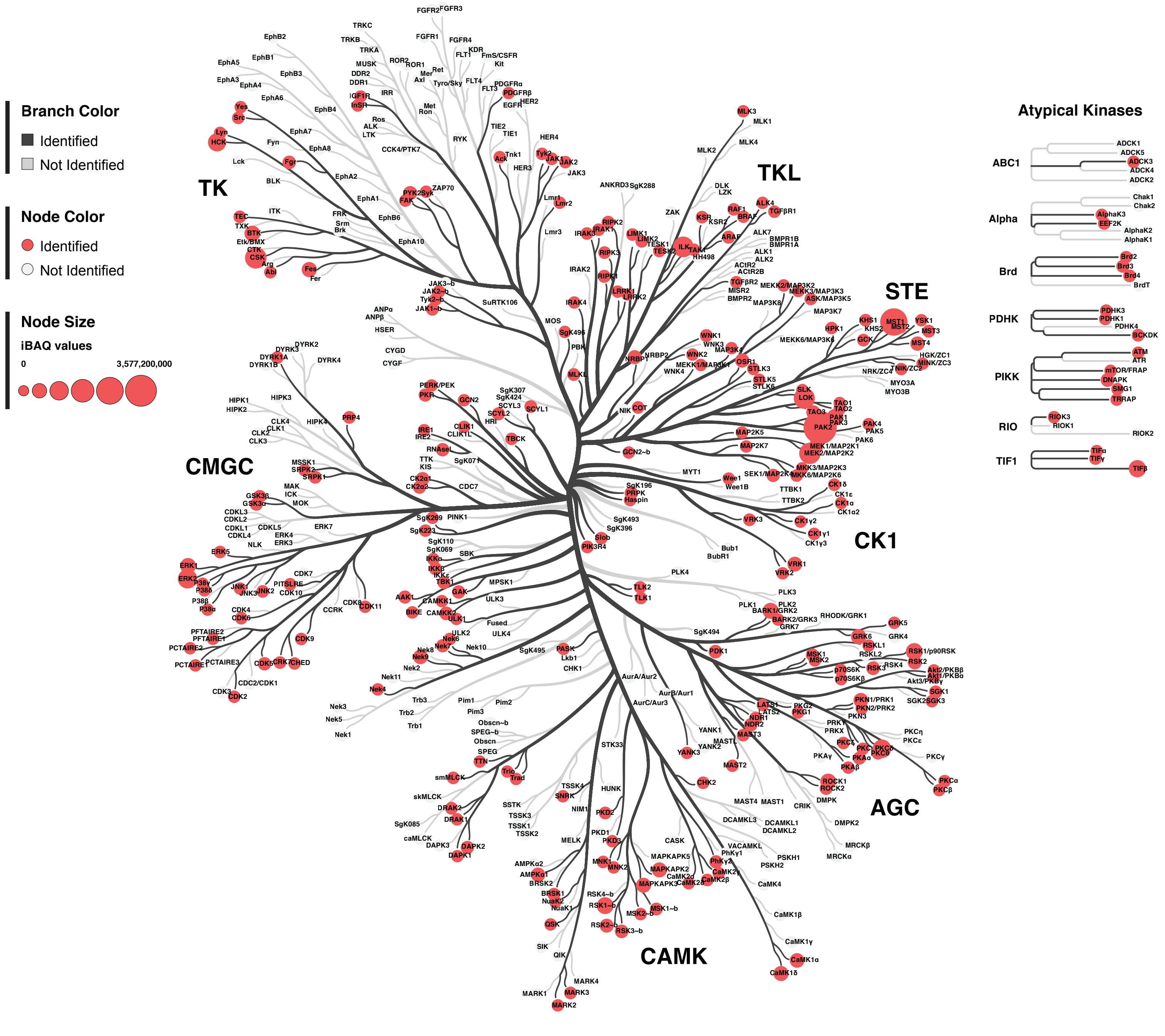

**Supplemental Figure 1**

(**A** and **B**) Volcano plots of phosphosites identified in the IL5- (A) and IL33- (B) non-activated eosinophil datasets. The $\log_2$ fold-change (x-axis) indicates the mean change in phosphorylation of each phosphosite. The $-\log_{10}$ q-value (y-axis) indicates the statistical significance of each changing phosphosite. Each dot represents one phosphosite. The grey dots represent no significant change between activated and non-activated eosinophils, the colored dots represent up-regulated (right) and down-regulated (left) phosphosites.

**Supplemental Figure 2**

(**A**) Volcano plot of phosphosites identified in the IL5-versus-IL33 datasets. The $\log_2$ fold-change (x-axis) indicates the mean change in phosphorylation of each phosphosite. The $-\log_{10}$ q-value (y-axis) indicates the statistical significance of each changing phosphosite. Each dot represents one phosphosite. The grey dots represent no significant change between the two activated states, the orange dots represent up-regulated (left) phosphosites in IL5 treatment and blue dots represent up-regulated (right) phosphosites in IL33 treatment. The top changing phosphosites were labeled by protein name. (**B**) Rank-order plot of phosphosites that are enriched in the IL5-versus-IL33 datasets. The phosphosites were ordered (0-16,691 on the x-axis) from lowest to highest fold change (y-axis). Each dot represents one phosphosite. The grey dots represent no significant change between the two activated states, the orange dots represent up-regulated (left) phosphosites in IL5 treatment and blue dots represent up-regulated (right) phosphosites in IL33 treatment. The top changing phosphosites were labeled by protein name.

**Supplemental Figure 3**

Kinome analysis from unstimulated human eosinophil proteomics datase[1]. A "Tree" plot generated by Coral[2] in which identified kinases are encoded in branch color and node color, and quantification (iBAQ) values are encoded in node size. Kinase families are noted in bold, with individual kinase nodes at the end of each branch.

#### SUPPLEMENTAL METHODS

**Purification of eosinophils.** Human blood eosinophils were purified from volunteer subject as described previously[3]. The study was approved by the University of Wisconsin-Madison Health Sciences Institutional Review Board, and informed written consent was obtained. Subjects were from a large volunteer pool of individuals with allergic rhinitis or non-severe asthma (**Supplementary Table 1**).

**Activation and preparation of eosinophils for proteomic and microscopic analysis.** On the same day as the blood donation and purification, eosinophils were received on ice in RPMI with 10% bovine serum. Cells were collected by centrifugation, resuspended in RPMI with 0.1% albumin, and incubated for 1 h at 37°C prior to further experimentation.

For proteomics, the cells were divided into 2 polypropylene tubes, each containing approximately 10 million cells in 10 ml. Tubes were incubated for 10 min with IL33, 50 ng/ml, or medium alone; IL5, 50 ng/ml, or medium alone; or IL33, 50 ng/ml, or IL5, 50 ng/ml. Cells were collected by a 1-min centrifugation, medium was removed by aspiration, and tubes with their cell pellets were immersed in liquid nitrogen and stored at -80°C.

When sets of samples from five individuals subjected to the same treatment had been collected, samples were thawed and compared by TMT analysis as described in the Supplement.

Immunofluorescence was performed on cells treated for 10 min with IL33, 50 ng/ml; IL5, 50 ng/ml; or medium alone in 4-ml polystyrene tubes as described previously[4]. As described in detail in the figure legend, cells with fixed with paraformaldehyde, collected by centrifugation, resuspended, cytospun onto coverslips, permeabilized, and stained for DNA and for antigens by double immuno-fluorescence using antibodies described in **Supplementary Table 2.**

**Comparative quantitation of protein and phosphopeptide abundances.** All steps were performed for three independent sets of samples (IL33 versus PBS, IL5 versus PBS, and IL33 versus IL5) (**Figure 1A**). The samples were thawed, resuspended in 6 M guanidine HCl (pH 8), and boiled at 100°C for 5 minutes. Methods are described in detail elsewhere[5] for generation of tryptic peptides, isobaric labeling (TMT 10-plex, Thermo Scientific), phosphopeptide enrichment, chromatographic separation of non-enriched and enriched peptides into pools by high pH reverse phase chromatography, and liquid chromatography MS/MS analysis of non-enriched peptide and phosphopeptide pools. Two technical replicates were analyzed for each phosphopeptide fraction, and data were combined to increase the yield of identified peptides. Raw data were processed with MaxQuant (version 1.5.2.8)[6,7]. Searches were performed against a target-decoy database of reviewed proteins plus isoforms (downloaded from UniProt). Recommended settings were used, and match-between-runs was not used. Cysteine carbamidomethylation and methionine oxidation were set to fixed and variable modifications, respectively. Phospho(STY) was added as a variable modification for the phosphopeptide searches. Phosphosites with a localization probability greater than 0.75 (Class I sites) were considered localized. Further processing was performed with Perseus (version 1.6.7.0)[8]. Statistical significance (q-value) was determined using a pairwise t-test with the Benjamini-Hochberg p-value correction. Raw files have been deposited to the MassIVE database with the identifier MSV000086364. Reviewers can access the files from ftp://MSV000086364@massive.ucsd.edu with the password “eos.”

**Production and characterization of antibodies to RAB44 and NHSL2.** RAB44 (Uniprot: Q7Z6P3), RAB46 (EFC4B_Human; Uniprot: Q9BSW2-2), NHSL2 (Uniprot: Q5HYW2) cDNA segments were amplified from eosinophil RNA by RT-PCR. The cDNAs were used as template for PCR to amplify full-length RAB44 (M1 to S1021), RAB44 N-terminal calmodulin-repeat domain (M1 to S379), RAB44 proline-rich domain (P380 to A828), RAB46 N-terminal calmodulin-repeat domain (M1 to R373), RAB46-proline-rich domain (E374 to S540), and NHSL2 mid-protein (R385 to E486). The cDNAs were cloned into the bacterial expression vector pET-ELMER by standard molecular biology techniques[9]. Protein expression of the constructs was induced in BL21(DE3) cells. Proteins were extracted with 8M urea and purified by immobilized metal affinity chromatography followed by ion-exchange chromatography as described previously for other constructs[10]. Rabbit anti-RAB44 proline-rich domain (WI610) and anti-NHSL2 mid-protein (W1612) polyclonal antibodies were produced through Covance Custom Antibody Development Services. The rabbits were immunized against 250 µg recombinant protein followed by three 125-µg booster injections. Antibodies were affinity-purified from the immune sera using the immunogen coupled to cyanogen bromide-activated Sepharose

4B (#C9142, Sigma-Aldrich) as described[10]. As non-immune controls, IgG was purified on Protein A Sepharose (P3391, Sigma-Aldrich) from serum obtained from each of the rabbits prior to immunization.

#### SUPPLEMENTARY TABLES

| Subject | Study | Condition | Age | Gender | Medications |
|---|---|---|---|---|---|
| 1 | IL33/PBS | Allergy no asthma | 25 | F | None |
| 2 | IL33/ PBS | Allergy no asthma | 37 | M | None |
| 3 | IL33/ PBS | Allergy no asthma | 45 | M | None |
| 4 | IL33/ PBS | Allergy and mild asthma | 22 | M | None |
| 5 | IL33/ PBS | Allergy and mild asthma | 31 | F | ICS/LABA |
| 6 | IL5/ PBS | Allergy no asthma | 21 | F | None |
| 7 | IL5/ PBS | Allergy and mild asthma | 36 | M | None |
| 8 | IL5/ PBS | Allergy and mild asthma | 45 | M | None |
| 9 | IL5/ PBS | Allergy no asthma | 51 | M | None |
| 10 | IL5/ PBS | Allergy no asthma | 31 | F | None |
| 11 | IL33/IL5 | Allergy no asthma | 51 | F | None |
| 12 | IL33/IL5 | Allergy and mild asthma | 38 | M | None |
| 13 | IL33/IL5 | Allergy and mild asthma | 43 | M | ICS |
| 14 | IL33/IL5 | Allergy and mild asthma | 34 | M | None |
| 15 | IL33/IL5 | Allergy and mild asthma | 27 | M | None |

**Supplementary Table 1: Characteristics of subjects whose eosinophils were studied.** ICS, inhaled corticosteroid. LABA, long-acting beta-agonist.

| Antibody | Source | Description | Catalog # | Concentration used |
|---|---|---|---|---|
| Anti-STAT3 (pY705) | Abcam | Rabbit mAb | ab76315 | 0.73 µg/ml |
| Anti-RELA | CST | Rabbit mAb | 8242S | 0.42 µg/ml |
| Anti-RAB46 | Proteintech | Mouse mAb | 66787-1-Ig | 2.0 µg/ml |
| Anti-RAB44 (WI610) | This study | Affinity-purified rabbit pAb | NA | 0.5 µg/ml |
| Anti-SRRM2 | ThermoFisher Scientific | Affinity-purified rabbit pAb | PA559559 | 1 µg/ml |
| Anti-SON | Santa Clara | Mouse mAb | sc-398508 | 1 µg/ml |

| | | | | |
|---|---|---|---|---|
| Anti-COIL | Santa Clara | Mouse mAb | sc-55594 | 1 µg/ml |
| Anti-NHSL2 (WI607) | Mosher lab | Affinity-purified rabbit pAb | NA | 1 µg/ml |
| Anti-VIM | Abcam | Mouse mAb | 8978 | 1 µg/ml |
| Anti-pP38 (pTGpY) | ThermoFisher Scientific | Mouse mAb | MA515218 | 2.4 µg/ml |
| Anti-pERK (pTEpY) | CST | Rabbit mAB | 4370S | 2.5 µg/ml |
| Anti-PML | Abcam | Rabbit mAb | 179466 | 2.5 µg/ml |
| Anti-TNFAIP3 | ThermoFisher Scientific | Mouse mAb | MA516164 | 2 µg/ml |
| Alexa488-labeled anti-rabbit IgG | ThermoFisher Scientific | Donkey pAb | A21206 | 4.0 µg/ml |
| Alexa555-labeled anti-mouse IgG | ThermoFisher Scientific | Donkey pAb | A31570 | 2.0 µg/ml |

**Supplementary Table 2: Sources and data on antibodies**.

| **Gene** | **Forward primer (5'-3')** | **Reverse primer ((5'-3')** |
|---|---|---|
| IL18 | TCTTCATTGACCAAGGAAATCGG | GGTCCGGGGTGCATTATCTCTA |
| CCL5 | TACACCAGTGGCAAGTGCTCC | CTCTGGGTTGGCACACACTTG |
| TNFSF14 | TGATACAAGAGCGAAGGTCTCACG | CTGAGTCTCCCATAACAGCGGC |
| CSF1 | CCTGGCGAGCAGGAGTATCAC | AGGTCTCCATCTGACTGTCAATCAG |
| PIM1 | GAGAGGCCCGACAGTTTCG | CTCCCCTTTCCGTGATGAAGT |
| BCL2 | GGTGGGGTCATGTGTGTGG | CGGTTCAGGTACTCAGTCATCC |
| CD69 | TGATGCCACCAGTCCCCAT | TCATTACAGCACACAGGACAGGTT |
| TP53 | CAGCACATGACGGAGGTTGT | TCATCCAAATACTCCACACGC |
| GAPDH | AGCCACATCGCTCAGACAC | GCCCAATACGACCAAATCC |

**Supplementary Table 3: List of assessed genes and the primers used by qPCR to measure mRNA at 2h activation.** Primer sequences were previously validated by the PrimerBank[11].

**Supplementary Spreadsheet 1.**

**Tab 1** contains a description of the data. The proteomics of IL5vNT (**Tab 2**), IL33vNT (**Tab 3**), and IL33vIL5 (**Tab 4**) are followed by the phosphoproteomics of IL5vNT (**Tab 5**), IL33vNT (**Tab 6**), and IL33vIL5 (**Tab 7**).


1. Wilkerson, E.M., Johansson, M.W., Hebert, A.S., Westphall, M.S., Mathur, S.K., Jarjour, N.N., Schwantes, E.A., Mosher, D.F., and Coon, J.J. (2016). The Peripheral Blood Eosinophil Proteome. J. Proteome Res. *15*, 1524–1533. 10.1021/acs.jproteome.6b00006.

2. Metz, K.S., Deoudes, E.M., Berginski, M.E., Jimenez-Ruiz, I., Aksoy, B.A., Hammerbacher, J., Gomez, S.M., and Phanstiel, D.H. (2018). Coral: Clear and Customizable Visualization of Human Kinome Data. Cell Syst. *7*, 347-350.e1. 10.1016/j.cels.2018.07.001.

3. Barretto, K.T., Swanson, C.M., Nguyen, C.L., Annis, D.S., Esnault, S.J., Mosher, D.F., and Johansson, M.W. (2018). Control of cytokine-driven eosinophil migratory behavior by TGF-beta-induced protein (TGFBI) and periostin. PLOS ONE *13*, e0201320. 10.1371/journal.pone.0201320.

4. Han, S.-T., and Mosher, D.F. (2013). IL-5 Induces Suspended Eosinophils to Undergo Unique Global Reorganization Associated with Priming. Am. J. Respir. Cell Mol. Biol. *50*, 654–664. 10.1165/rcmb.2013-0181OC.

5. Muehlbauer, L.K., Wei, T., Shishkova, E., Coon, J.J., and Lambert, P.F. (2022). IQGAP1 and RNA Splicing in the Context of Head and Neck via Phosphoproteomics. J. Proteome Res. *21*, 2211–2223. 10.1021/acs.jproteome.2c00309.

6. Cox, J., and Mann, M. (2008). MaxQuant enables high peptide identification rates, individualized p.p.b.-range mass accuracies and proteome-wide protein quantification. Nat. Biotechnol. *26*, 1367–1372. 10.1038/nbt.1511.

7. Cox, J., Neuhauser, N., Michalski, A., Scheltema, R.A., Olsen, J.V., and Mann, M. (2011). Andromeda: A Peptide Search Engine Integrated into the MaxQuant Environment. J. Proteome Res. *10*, 1794–1805. 10.1021/pr101065j.

8. Tyanova, S., Temu, T., Sinitcyn, P., Carlson, A., Hein, M.Y., Geiger, T., Mann, M., and Cox, J. (2016). The Perseus computational platform for comprehensive analysis of (prote)omics data. Nat. Methods *13*, 731–740. 10.1038/nmeth.3901.

9. Maurer, L.M., Tomasini-Johansson, B.R., Ma, W., Annis, D.S., Eickstaedt, N.L., Ensenberger, M.G., Satyshur, K.A., and Mosher, D.F. (2010). Extended Binding Site on Fibronectin for the Functional Upstream Domain of Protein F1 of Streptococcus pyogenes. J. Biol. Chem. *285*, 41087–41099. 10.1074/jbc.M110.153692.

10. Bortnov, V., Tonelli, M., Lee, W., Lin, Z., Annis, D.S., Demerdash, O.N., Bateman, A., Mitchell, J.C., Ge, Y., Markley, J.L., et al. (2019). Solution structure of human myeloid-derived growth factor suggests a conserved function in the endoplasmic reticulum. Nat. Commun. *10*, 5612. 10.1038/s41467-019-13577-5.

11. PrimerBank: a PCR primer database for quantitative gene expression analysis, 2012 update | Nucleic Acids Research | Oxford Academic https://academic.oup.com/nar/article/40/D1/D1144/2902573.
